# Mitocurcumin mediated redox disruption and metabolic rewiring induces tumor regression in *Drosophila* intestinal stem cell tumors

**DOI:** 10.64898/2026.08.27.747455

**Authors:** Somesh Diwate, Ujjayita Chowdhury, Nikhil Gadewal, Shraddha Jadhav, Vikram Gota, Rohan Jayant Khadilkar

## Abstract

Mitochondria-targeted modulation of redox homeostasis has emerged as a promising strategy for controlling pathological cell proliferation. Here, we investigate the effects of Mitocurcumin in a Yorkie-driven intestinal stem cell tumor model in *Drosophila.* Using an integrative, genetically tractable approach combining in silico molecular modelling with in vivo functional analyses, we identify thioredoxin reductase (TrxR) as a conserved redox-associated target of Mitocurcumin. Docking and molecular dynamics simulations predict a stable interaction of Mitocurcumin with both *Drosophila* and mammalian TrxR homologs. Functionally, Mitocurcumin treatment reduces mitotic activity, elevates reactive oxygen species (ROS) selectively within escargot-positive intestinal stem cell population, enhances apoptosis in the tumor-bearing guts, and causes significant mitochondrial membrane depolarization. These cellular effects coincide with dose-dependent regression of Yorkie-induced intestinal hyperplasia. Despite mitochondrial functional impairment, mitochondrial morphology remains largely preserved, suggesting primary disruption of redox buffering rather than structural collapse. Metabolomic profiling of these guts further reveals remodelling of energy metabolism consistent with adaptive responses to oxidative stress. Importantly, Mitocurcumin alleviates tumor-associated organismal bloating and significantly extends lifespan indicating a previously uncharacterized systemic, organism-wide response to Mitocurcumin treatment in an in vivo scenario. Collectively, our findings establish TrxR-mediated redox regulation as a critical vulnerability in Yorkie-driven hyperproliferation and highlight the utility of *Drosophila* as an integrative in vivo platform for evaluating mitochondria-targeted bioactive molecules.

## Introduction

Colorectal cancer (CRC) remains a major cause of cancer-related mortality worldwide, as reflected by updated GLOBOCAN 2022 statistics emphasizing its substantial global impact (Bray et al., 2024). In CRC, the transcriptional co-activators, Yes-associated protein (YAP) and Transcriptional co-activator with PDZ-binding motif (TAZ) are frequently overactivated, and their nuclear accumulation is strongly linked to aggressive clinical outcomes by promoting proliferation, stem-like features, and therapy resistance(Zanconato et al., 2016). Classically known to regulate cell division, emerging studies suggest oncogenic YAP has extensive roles in reshaping cellular metabolism. YAP/TAZ reprograms metabolism by boosting glycolysis, enhancing glutamine/amino acid uptake and driving lipid accumulation in liver(Koo & Guan, 2018; Noto et al., 2017). To cope with the elevated metabolic demands, cancer cells must also counteract oxidative stress resulting from increased reactive oxygen species (ROS). YAP contributes to this adaptation by enhancing antioxidant defences and maintaining redox balance favourable for tumor survival(Dai et al., 2022).

The dependence of YAP-driven tumors on mitochondrial and redox homeostasis highlights a potential therapeutic vulnerability. Given the pivotal role of mitochondria in bioenergetics and signaling, their metabolism has emerged as a promising target for cancer therapy(Weinberg & Chandel, 2015). The thioredoxin (Trx) system, a key antioxidant defense mechanism, regulates redox equilibrium, prevents oxidative damage, and supports proliferative signaling, making it an attractive candidate for therapeutic intervention(Seitz et al., 2024). In tumors, elevated TrxR (Thioredoxin reductase) levels help cancer cells tolerate high ROS levels due to increased proliferation and metabolic alterations supporting survival, sustained proliferation, invasion and therapy resistance(Karlenius & Tonissen, 2010; Muri & Kopf, 2023; Shan et al., 2010). Targeting TrxR could be highly effective as it would disrupt the ROS balance thereby inducing apoptosis. Some examples of candidate molecules that have shown efficacy and immense potential are auranofin (targets TrxR selenocysteine), PX-12 (binds Trx), and selenium-based agents like EbD, which form covalent Se-Se/Se-S bonds with TrxR’s active site for irreversible inhibition. These have been shown to have preclinical efficacy in lung, breast, and other cancers(Bian et al., 2019; Pan et al., 2020; Tang et al., 2024). However, the in vivo metabolic and redox consequences of inhibiting this system in YAP-dependent tumors remain poorly understood.

Mitocurcumin, a mitochondria-targeted derivative of curcumin, exhibits superior cellular uptake and bioavailability compared with native curcumin (Reddy et al., 2014). Conjugation with a triphenylphosphonium (TPP) moiety enables its mitochondrial accumulation, where it inhibits TrxR2 activity, elevates ROS levels, activates JNK signaling, and induces cancer cell death (Gaur et al., 2024; Jayakumar et al., 2017). Reports have also suggested MitoC induces paraptosis-like cell death in lung cancer cells by triggering ROS-mediated JNK signaling offering a potential strategy to overcome apoptosis-resistant chemoresistance in lung cancer (Panigrahi et al., 2026). Genetic conservation, low genetic redundancy, sophisticated gene manipulation and short life cycle of *Drosophila* provide an advantage for modelling human tumors as well as whole-organism anticancer drug screening (Cagan et al., 2019). It has been estimated that approximately 75% of disease-causing genes in humans have functional homologs in *Drosophila*(Wangler et al., 2017). While mammals possess two thioredoxin reductases, *Drosophila* primarily relies on a cytosolic form, TrxR1, with limited mitochondrial specialization(Kanzok et al., 2001; Missirlis et al., 2002). *Drosophila* intestine provides a robust model for investigating tumor biology and drug responses due to its conserved signaling pathways and well-characterized stem cell compartment(Jiang & Edgar, 2012; Micchelli & Perrimon, 2006). The Hippo pathway, evolutionarily conserved from flies to mammals, operates through the transcriptional co-activator Yorkie (Yki), the *Drosophila* homolog of mammalian YAP(Harvey et al., 2003). Targeted overexpression of constitutively active Yki in the intestinal epithelium effectively models oncogenic transformation, establishing the *Drosophila* midgut as a genetically tractable system for studying Hippo pathway–driven tumorigenesis and for in vivo drug screening(Bajpai et al., 2020; Karpowicz et al., 2010; Shaw et al., 2010). The adult midgut tumor recapitulates not just tumor-autonomous effects but also systemic effects like organ wasting and bloating seen typically in cancer-associated cachexia(Figueroa-Clarevega & Bilder, 2015; Hsi et al., 2023; Kwon et al., 2015). These hyperplastic ISC tumors in the adult midgut are well positioned to mimic the complex interplay between local and systemic effects, making them a widely accepted tumor model for studying therapeutics.

In this study, we demonstrate that mitochondrial redox modulation by Mitocurcumin suppresses Yorkie-driven stem cell tumor load in the *Drosophila* midgut. By integrating in-silico analyses of TrxR interaction with mitocurcumin along with in-vivo functional assays to investigate its effect on the tumor burden and metabolism, we identify thioredoxin reductase (TrxR) as a central redox node linking mitochondrial dysfunction to tumor regression in a Hippo/Yorkie-dependent context. Our study provides significant insights on how targeting the finely tuned ROS balance could be a promising therapeutic modality for tackling tumors.

## Results

### In-silico validation shows mitocurcumin binding to thioredoxin reductase

Mitocurcumin, a mitochondria-targeted derivative generated by conjugation of curcumin to the lipophilic triphenylphosphonium (TPP) cation, has been shown to exhibit enhanced stability and improved biological efficacy compared to native curcumin (Supplementary Fig. S1A-B) (Reddy et al., 2014). Although curcumin is a natural compound with well-documented anti-inflammatory and anti-cancer properties, its clinical utility is limited because of poor bioavailability and stability in clinical settings. Previous reports suggest Mitocurcumin can inhibit Thioredoxin Reductase (TrxR) activity, supported by in silico docking analyses and enzymatic inhibition assays(Jayakumar et al., 2017). In the present study, we aimed to determine whether Mitocurcumin also exhibits favorable binding to *Drosophila* TrxR. Thioredoxin reductase (TrxR) exists as functionally conserved isoforms across species, despite differences in their C-terminal redox-active residues. In *Drosophila*, TrxR lacks the canonical C-terminal selenocysteine (Sec) residue present in mammalian TrxRs and instead utilizes a cysteine residue while retaining high catalytic activity (Missirlis et al., 2002).To evaluate whether Mitocurcumin can directly associate with *Drosophila* TrxR, comparative in silico analyses were performed using *Drosophila* TrxR and mammalian TrxR. Multiple sequence alignment of chain A revealed strong conservation within the catalytic core and cofactor-binding regions (Supplementary Fig. S1C), and 3D structural superposition confirmed conservation of overall fold and active-site architecture where Mitocurcumin binds (Fig. 1A). Molecular docking revealed favorable binding of Mitocurcumin to both enzymes. In *Drosophila* TrxR1, Mitocurcumin exhibited a Glide score of −9.02 kcal/mol, while the mammalian homolog yielded a score of −8.37 kcal/mol (Table 1). In both cases, Mitocurcumin occupied a comparable binding pocket near the FAD-binding site (Fig. 1B-C). Detailed protein–ligand interaction analysis showed that Mitocurcumin engages a defined network of residues (Table 1). In *Drosophila* TrxR1, the complex is stabilized by a key π–π stacking interaction with Tyr197 and hydrogen bonds with Lys65 and Glu333. In the mammalian complex, a similar anchor is formed via π–π stacking with Tyr200, supported by interactions with Lys68, Lys69, and Ser199. These aromatic interactions with conserved Tyrosine residues are strategically positioned near the redox-active centres, suggesting that Mitocurcumin may interfere with the electron transfer process required for catalytic turnover. To assess the dynamic stability of the Mitocurcumin–TrxR complexes, molecular dynamics simulations of 500 nanoseconds were performed. RMSD analysis demonstrated rapid equilibration followed by stable fluctuations for both *Drosophila* TrxR1–Mitocurcumin and mammalian TrxR1–Mitocurcumin complexes (Fig. 1D-E). Ligand RMSD profiles further confirmed that Mitocurcumin remained stably bound within the pocket over time (Supplementary Fig. S1D–E). RMSF analysis revealed limited flexibility among residues constituting the ligand-binding pocket (Supplementary Fig. S1F–G). Contact frequency analysis throughout the trajectory highlighted Tyr197 as a critical anchor in *Drosophila* TrxR1, maintaining π–π stacking for over 80% of the simulation. In the mammalian complex, Lys69, Ser199, and Tyr200 served as the primary stabilizing residues (Fig. 1F-I). Binding free-energy estimation using MM/GBSA (Molecular Mechanics/Generalized Born Surface Area) yielded an average ΔG(NS) of −28.92 kcal/mol for the *Drosophila* TrxR1– Mitocurcumin complex and −6.51 kcal/mol for the mammalian TrxR–Mitocurcumin complex (Table 1), indicating energetically favorable binding in both systems.

**Figure 1.**
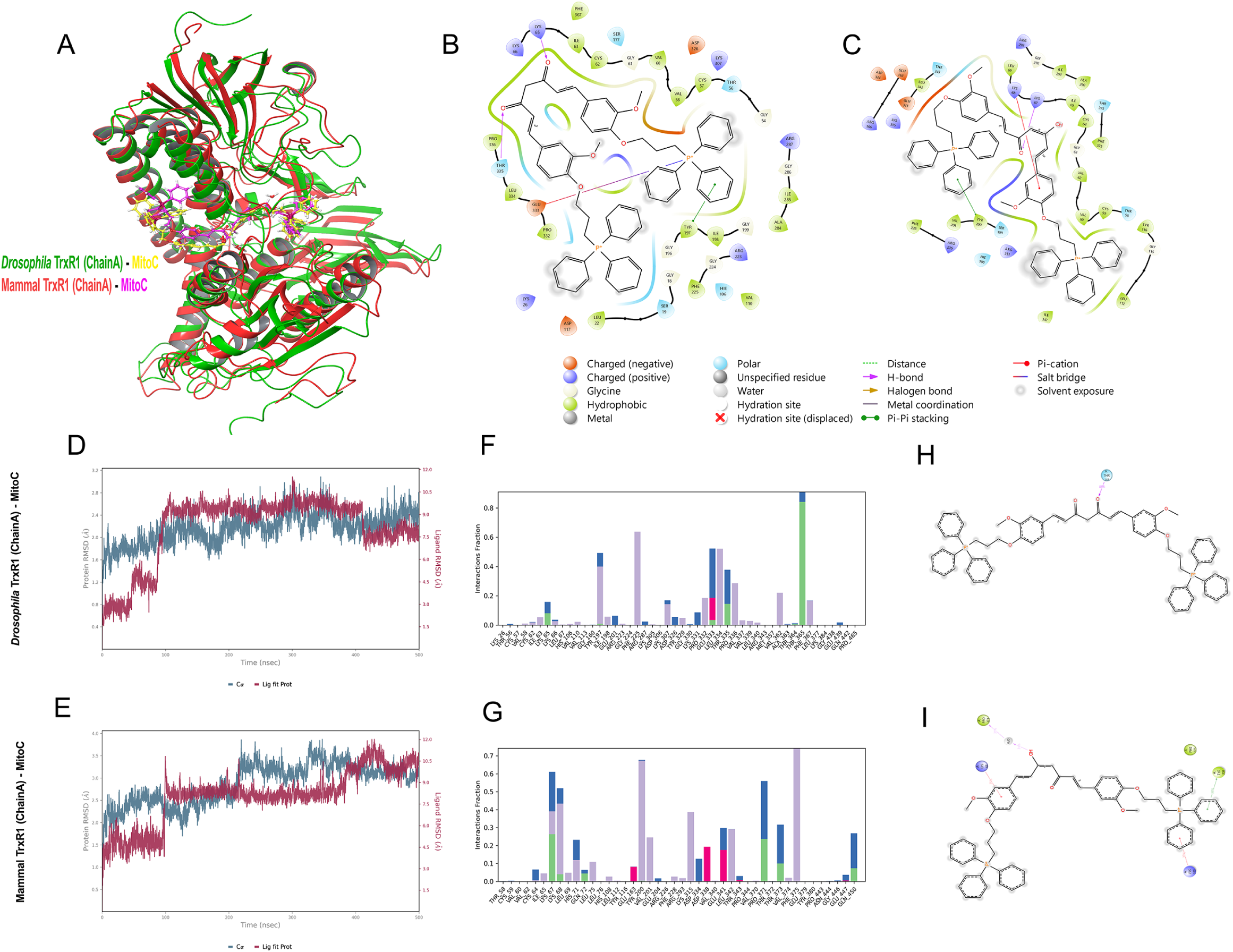
In silico characterization of Mitocurcumin binding to thioredoxin reductase (TrxR) Structural superposition of *Drosophila melanogaster* TrxR1 (Chain A; PDB: 3DH9) and mammalian TrxR (Chain A; PDB: 3QFA), highlighting conservation of the overall fold and FAD/NADPH-binding catalytic pocket. Structures are shown in ribbon representation (green, Drosophila; red, mammalian) (A). Two-dimensional protein–ligand interaction maps showing binding of Mitocurcumin (MitoC) within the catalytic site of Drosophila TrxR1 (B) and mammalian TrxR (C), including hydrogen bonding, hydrophobic, π–π stacking, and polar interactions. Backbone RMSD plots from 500 ns molecular dynamics simulations demonstrating structural stability of Mitocurcumin-bound complexes (D–E). Residue-wise interaction frequency plots of ligand–protein contacts throughout the simulation (F–I). Docking was performed using Glide (SP mode), and molecular dynamics simulations were conducted using Desmond with the OPLS4 force field under explicit solvent conditions.

This suggests that while the initial docking poses are similar, the *Drosophila* enzyme provides a more energetically favorable and stable environment for Mitocurcumin over time. The stronger energetic profile in *Drosophila* likely stems from a more robust hydrogen-bonding network and the greater stability of π-stacking interactions. Furthermore, because *Drosophila* TrxR1 utilizes cysteine residues instead of the mammalian selenocysteine residue, these results demonstrate that Mitocurcumin’s inhibitory mechanism is likely directed towards the conserved catalytic core and is independent of selenocysteine, supporting its utility as a broad-spectrum TrxR inhibitor. Together, these analyses demonstrate that *Drosophila* TrxR1 provides a structurally and dynamically compatible binding environment for Mitocurcumin, comparable to that of mammalian TrxRs, supporting the relevance of the *Drosophila* model for investigating TrxR-associated redox phenotypes.

### Mitocurcumin drives dose-dependent regression of activated Yorkie induced intestinal tumors

The *Drosophila* adult midgut consists of an epithelium that is maintained by the multipotent population of intestinal stem cells (ISCs) that either regenerate the stem cell pool or differentiate to maintain intestinal homeostasis(Lemaitre & Miguel-Aliaga, 2013; Miguel-Aliaga et al., 2018). Hippo signaling pathway is a conserved regulator of tissue growth and homeostasis, with Yorkie (Yki) acting as its transcriptional co-activator in *Drosophila* (Harvey et al., 2003; Kango-Singh et al., 2002). Activating mutations in the Yki proto-oncogene in ISCs result in robust hyperplasia and an effective tumor model in *Drosophila*(Kwon et al., 2015; Oh & Irvine, 2008; Oh & Irvine, 2009). We incubated the freshly eclosed adult flies bearing the Yki induced tumors at 18 °C for 3 days after which we began induction of the Gal4 at 29 °C for 48h. Tumor bearing flies were treated with either Mitocurcumin or DMSO during this 48 hours induction period and the tumor load was assessed after the 48h time window (Fig. 2A). We observe that there is a dose-dependent reduction in the Esg-GFP^+^ positive tumor load upon Mitocurcumin administration (Fig. 2B-L). We sought out to identify a mitocurcumin concentration capable of reducing the Esg⁺ tumor burden by approximately 50%. To establish a quantitative benchmark, we used the Esg⁺ cell load observed after 24h of Yki induction as a reference, as this represented roughly half of the Esg⁺ expansion seen at 48h (Fig. 2L). We therefore aimed to determine the Mitocurcumin concentration that could reduce the 48h Esg⁺ load to levels comparable to the 24h induction condition (Fig. 2B–C). Mitocurcumin treatment elicited a graded, concentration-dependent reduction in tumor burden. While 10 µM and 20 µM concentration of Mitocurcumin resulted in a partial reduction of Esg-positive cell clusters, treatment with 30 µM resulted in robust tumor reduction accompanied by partial rescue of gut architecture (Fig. 2E–I). Increasing the concentration to 40 µM did not lead to further suppression of tumor growth, indicating a plateau in therapeutic efficacy (Fig. 2J–L). Based on these observations, 30 µM Mitocurcumin was selected as the optimal dose for subsequent experiments.

**Figure 2.**
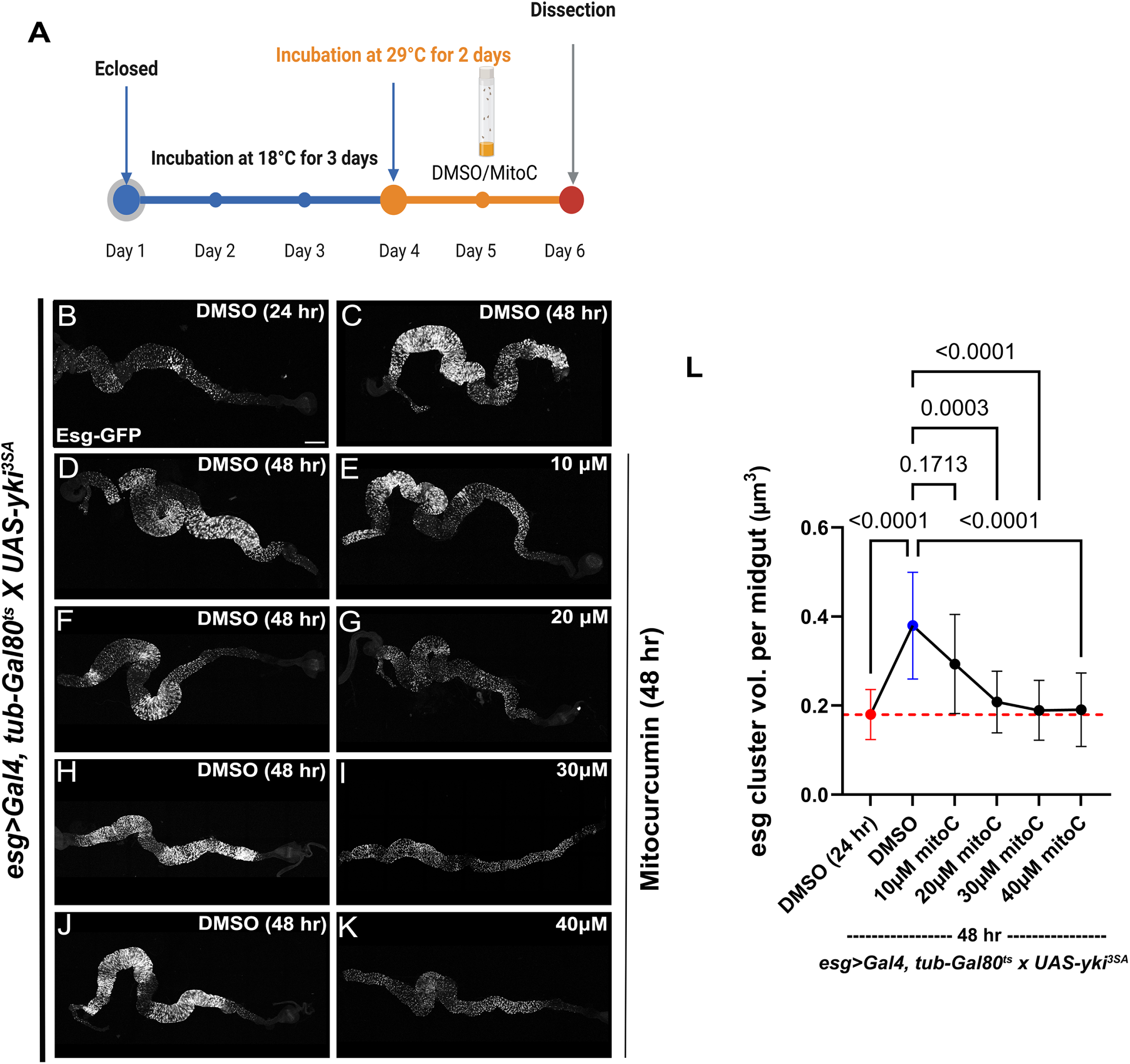
Mitocurcumin drives dose-dependent regression of activated Yorkie-induced intestinal tumors. Experimental design for tumor induction and drug treatment in *esg-GAL4, tub-GAL80^ts^ > UAS-yki^3SA^* flies (A). Whole mount adult gut displaying esg-positive cells from flies treated with increasing concentrations of Mitocurcumin; 10 µM (E), 20 µM (G), 30 µM (I), 40 µM (K) for 48 h compared to DMSO-treated midguts at 24h (B) and respective 48h post-induction (C, D, F, H, J).Quantification of tumor burden expressed as the ratio of esg-positive (GFP+) volume to total gut (L). Esg-GFP (Gray) marks the intestinal stem cells (ISCs). Data represents mean ± SEM (n = 9 midguts for DMSO 24h and 10 µM groups; n = 10 midguts for all other conditions). Statistical analysis was performed using one-way ANOVA with multiple comparisons. p-values indicating statistical significance have been mentioned in the respective graphs. Scale bar: 200 µm (B-K). Created in BioRender. Khadilkar, R. (2026) https://BioRender.com/ee340g3.

### Mitocurcumin limits proliferation and enhances apoptosis in Yorkie-driven tumors

To determine whether mitocurcumin-induced tumor reduction was associated with altered cellular proliferation or survival, midguts were analyzed after 48h of Yki induction under DMSO or mitocurcumin treatment. We observed pronounced hyperproliferation in DMSO-treated controls, as evidenced by a marked increase in phospho-Histone H3 (H3P)-positive mitotic cells within esg⁺ tumor regions (Fig. 3A–A″). Upon mitocurcumin treatment, the number of H3P-positive cells were significantly reduced (Fig. 3B–B″). Quantitative analysis revealed that Mitocurcumin decreased mitotic activity to levels comparable to those observed after 24h of Yki induction (Fig. 3C), indicating that mitocurcumin effectively suppresses hyperproliferation and restores proliferative activity to a control state in the *yki3SA* tumor-bearing flies.

**Figure 3.**
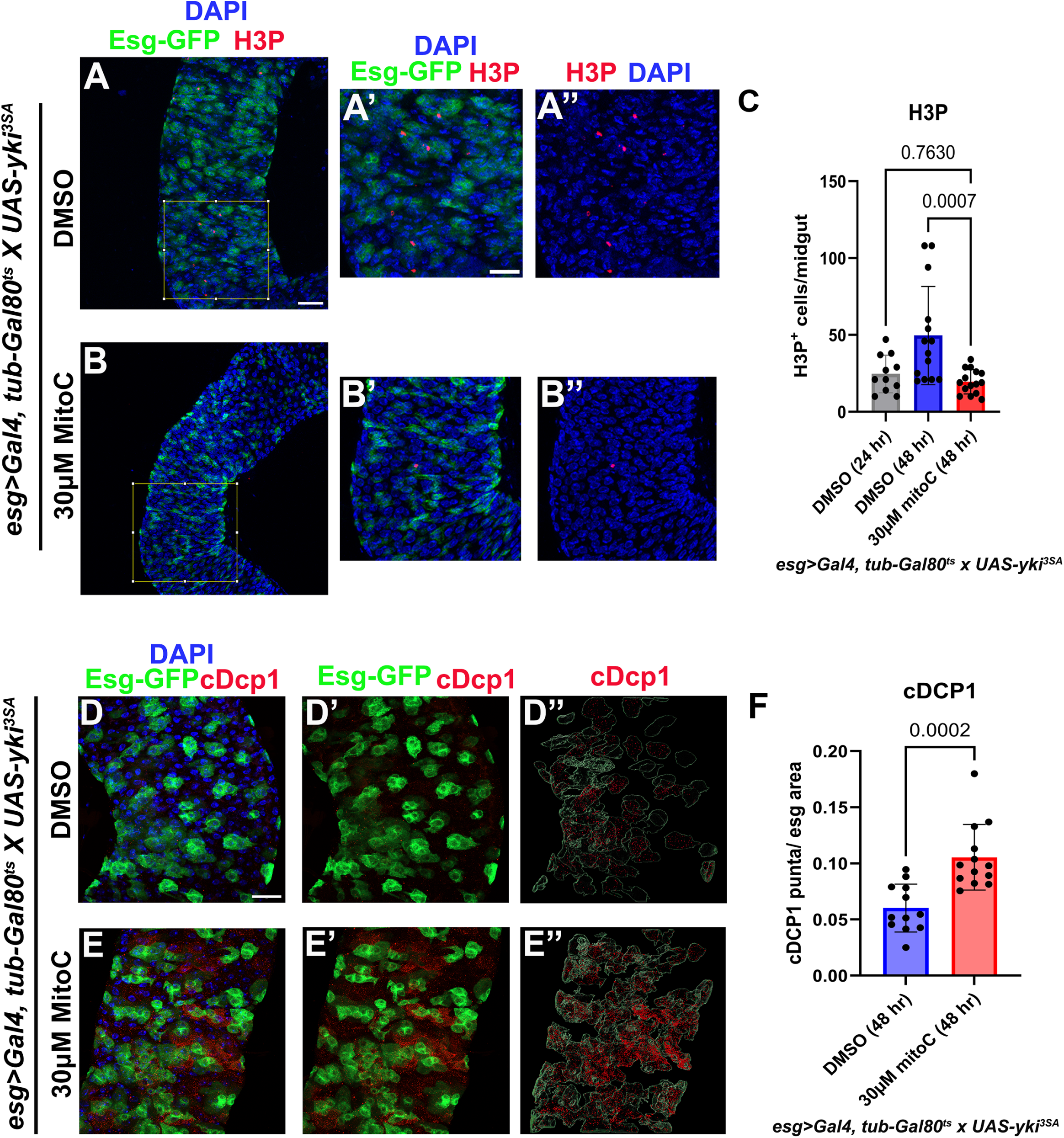
Mitocurcumin limits proliferation and enhances apoptosis in Yorkie-driven tumors. Adult midguts from tumor bearing flies stained with phospho-Histone H3 (pH3, Red) upon 30µM of Mitocurcumin treatment (B, B’, B’’) compared to DMSO treatment (A, A’, A’’) for 48h. Corresponding graph of quantification of H3P-positive cells per midgut for the respective groups (C). Adult midguts from tumor bearing flies stained with cleaved-Dcp1 (cDcp1, Red) upon 30 µM of Mitocurcumin treatment (E, E’, E’’) compared to DMSO treatment (D, D’, D’’) for 48h. Corresponding graph of quantification of cDcp1 puncta per esg cluster area for the respective groups (F). Data represent mean ± SEM (n = 10 midguts per group). Nuclei were stained with DAPI (Blue). Esg-GFP (Green) marks the intestinal stem cells (ISCs). D’’ and E’’ indicate segmented area in the midgut for cDcp1 quantification. Statistical significance was determined using Student’s t-test (F) or one-way ANOVA (C). p-values indicating statistical significance have been mentioned in the respective graphs. Scale bar: 30 µm (A, B, D, E) and 15 µm (A’, A’’, B’, B’’, D’, D’’, E’, E’’).

Given the reduction in Esg⁺ tumor burden (Fig. 2), we next investigated whether Mitocurcumin treatment promotes apoptosis within the tumor compartment. Cleaved *Drosophila* Caspase 1 (cDCP1) staining revealed minimal apoptotic signal in esg⁺ regions of DMSO-treated *yki^3SA^* tumors (Fig. 3D–D″). In contrast, Mitocurcumin-treated midguts displayed an accumulation of cDCP1-positive puncta within esg⁺ tumor regions (Fig. 3E–E″). To specifically quantify apoptosis within the tumor compartment, cDCP1-positive signals were segmented and measured within the Esg⁺ region. This analysis demonstrated a significant increase in apoptotic signals in Mitocurcumin-treated flies compared to control flies harbouring Yki-driven ISC tumors (Fig. 3F). Collectively, these findings indicate that mitocurcumin suppresses Yki-driven tumor expansion through a dual mechanism involving inhibition of mitotic activity and induction of apoptosis within the Esg-positive tumor compartment.

### Mitocurcumin increases ROS levels and activates antioxidant signaling in the ISC tumors in the midgut

ROS is maintained in an equilibrium in cancer cells wherein cancer cells upregulate antioxidant systems like SODs, catalase, glutathione, thioredoxin, and NRF2-driven detoxification programs to keep the excess ROS generated by cancer cells in check(Brandl et al., 2025). The shift in this equilibrium disturbs this fine balance leading to oxidative damage that can result in stress and apoptosis of cancer cells. In the *Drosophila* adult midgut, low to moderate ROS levels promote ISC proliferation and stem cell regeneration, whereas excessive ROS triggers apoptosis, differentiation defects, and cytotoxic effects(Chen et al., 2021). Comparative studies in *Drosophila* and mammalian systems highlight a conserved role for ROS in balancing ISC self-renewal versus cell death(Morris & Jasper, 2021). Having established that mitocurcumin suppresses mitotic activity and promotes apoptosis in Yki-driven tumors (Fig. 3), we next investigated whether these effects were associated with disruption of redox homeostasis within the tumor compartment.

To investigate whether Mitocurcumin perturbs redox balance in Yki-driven gut tumors, intracellular reactive oxygen species (ROS) were quantified. DMSO-treated control midguts displayed low basal ROS signals within Esg-positive tumor clusters (Fig. 4A–A″). In contrast, Mitocurcumin treatment induced a strong elevation of ROS signal specifically within Esg+ cell clusters (Fig. 4B–B″). Paraquat treatment was included as a positive control and produced robust ROS accumulation (Fig. 4C–C″). Measurement of ROS intensity within Esg-positive regions confirmed a significant increase in ROS levels following Mitocurcumin treatment relative to DMSO controls, reaching levels comparable to those induced by paraquat (Fig. 4D). Given that excessive ROS can activate stress-responsive antioxidant pathways, we next examined the Keap1–Cnc signaling axis that regulates oxidative stress tolerance in *Drosophila* (Sykiotis & Bohmann, 2008). In *Drosophila*, Cap ‘n’ collar (Cnc), the functional homolog of mammalian Nrf2, is negatively regulated by Kelch-like ECH-associated protein 1 (Keap1) under basal conditions (Hochmuth et al., 2011; Sykiotis & Bohmann, 2008) (Fig 4E). Oxidative stress disrupts Keap1-mediated repression, permitting Cnc stabilization and transcriptional activation of antioxidant genes (Sykiotis & Bohmann, 2008). We observed significant induction of *cnc* gene expression in mitocurcumin-treated tumor bearing flies, accompanied by reduced *keap1* expression (Fig. 4F, G). These findings are consistent with activation of the Cnc-dependent antioxidant response in tumor tissue experiencing elevated oxidative stress.

**Figure 4.**
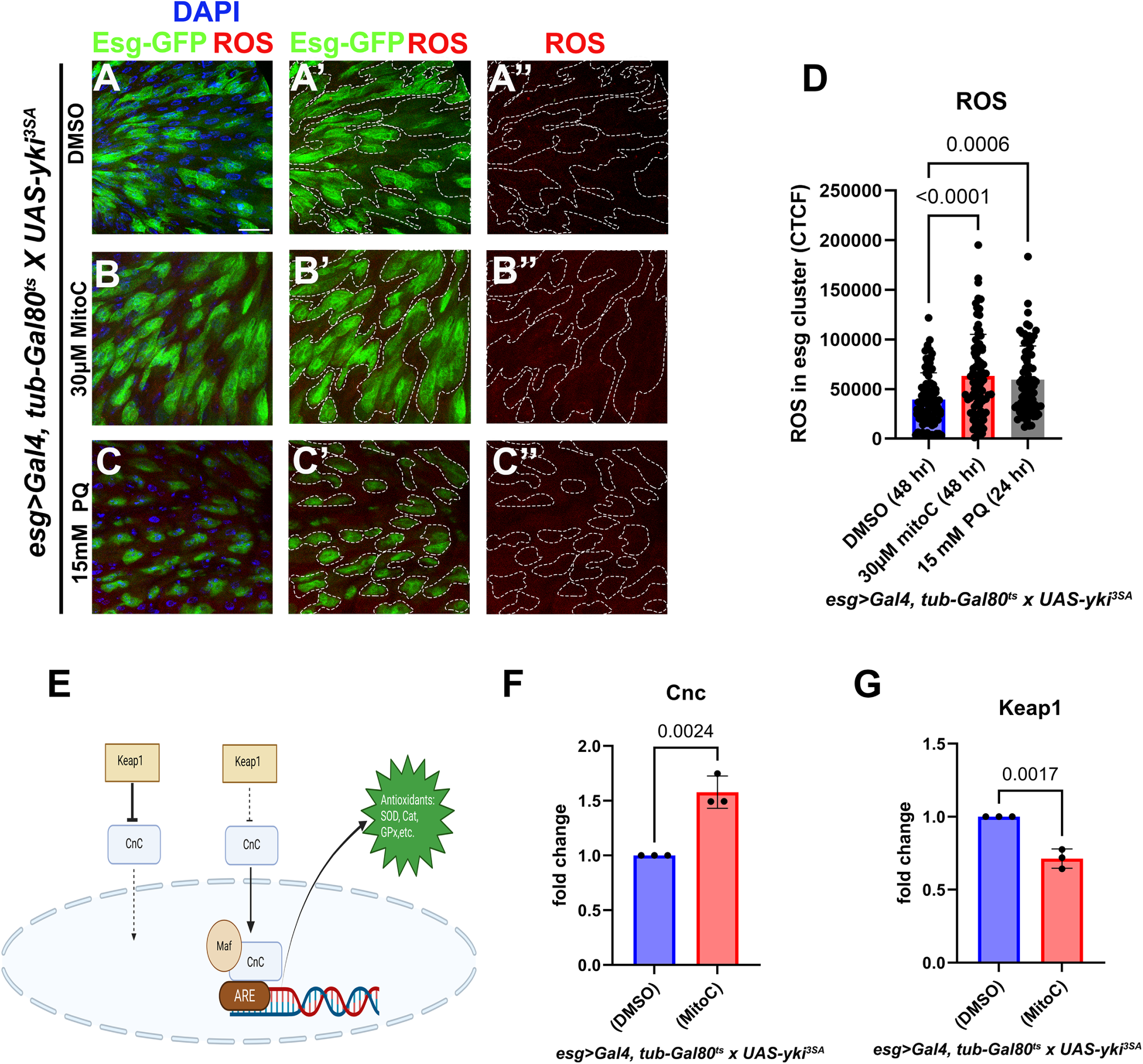
Mitocurcumin increases ROS levels and activates antioxidant signaling in the ISC tumors in the midgut. CellROX Deep Red (Red) assay for ROS detection in the midguts from tumor bearing flies flies to visualize ROS upon 30µM of Mitocurcumin treatment (B, B’, B’’) compared to DMSO treatment (A, A’, A’’) for 48h and 15 mM Paraquat (PQ) treatment (C, C’, C’’) for 24h. Corresponding graphs of quantification of ROS levels expressed as corrected total cell fluorescence (CTCF) within esg-positive regions (D). Dashed outlines indicate regions used for quantification. Schematic representation of the Cnc–Keap1–ARE antioxidant pathway (E). Graphs showing quantification of the relative mRNA expression of *cnc* and *keap1* (F, G). RNA was isolated from 10 midguts per biological replicate (n = 3 biological replicates), normalized to *rp49*. Nuclei were stained with DAPI (Blue in A, B, and C). Esg-GFP (Green) marks the intestinal stem cells (ISCs). Data represent mean ± SEM. Statistical significance was assessed using Student’s t-test. p-values indicating statistical significance have been mentioned in the respective graphs. Scale bar: 15 µm (A-C’’). Created in BioRender. Khadilkar, R. (2026) https://BioRender.com/ftd31da.

In order to assess and characterize the antioxidant enzymes that are modulated in the midgut tumors upon treatment with mitocurcumin, we used a similar 48h induction and treatment time window (Supplementary Fig. S2A) and observe that there is no significant change in *catalase* and *dSod2* mRNA levels whereas *gtpx* mRNA levels are downregulated by qPCR upon mitocurcumin treatment as compared to the DMSO control (Supplementary Fig. S2B-D). Taken together, these results indicate that mitocurcumin-induced tumor regression is associated with pronounced ROS accumulation and activation of compensatory antioxidant signaling within Esg⁺ tumor cells.

### Mitocurcumin does not significantly alter mitochondrial morphology in Yki-driven Esg⁺ tumor cells

Given that Mitocurcumin induced ROS accumulation (Fig. 4), we next examined whether this is accompanied by structural alterations in mitochondrial morphology within Esg⁺ tumor cells. To visualise mitochondria, immunostaining of the midguts was done with the MitoView probe. Our analysis revealed comparable mitochondrial organization in the EsgGFP+ cell clusters of DMSO- and Mitocurcumin-treated samples (Fig. 5A–B′′′). Mitochondria in both conditions displayed a reticular and tubular network distributed throughout the cytoplasm of EsgGFP⁺ cell clusters, without obvious fragmentation, swelling, or collapse. To quantitatively assess mitochondrial morphology, three-dimensional reconstruction and analysis were performed using the FIJI mitochondrial analyzer plugin. Multiple structural parameters were measured, including mitochondrial number per cell, number of branches, mean branch length, mean surface area, and mean volume. Quantitative analysis demonstrated no statistically significant differences between the mitochondria from DMSO- and Mitocurcumin-treated tumor bearing flies across all measured parameters (Fig. 5C–G). Specifically, mitochondrial number and branching complexity remained unchanged, and no significant differences were observed in mean branch length, surface area, or mitochondrial volume. Together, these data indicate that although Mitocurcumin specifically targets mitochondria and induces redox stress, it does not cause overt disruption of mitochondrial morphology within the 48h treatment window where we observe decrease in tumor load and apoptotic induction.

**Figure 5.**
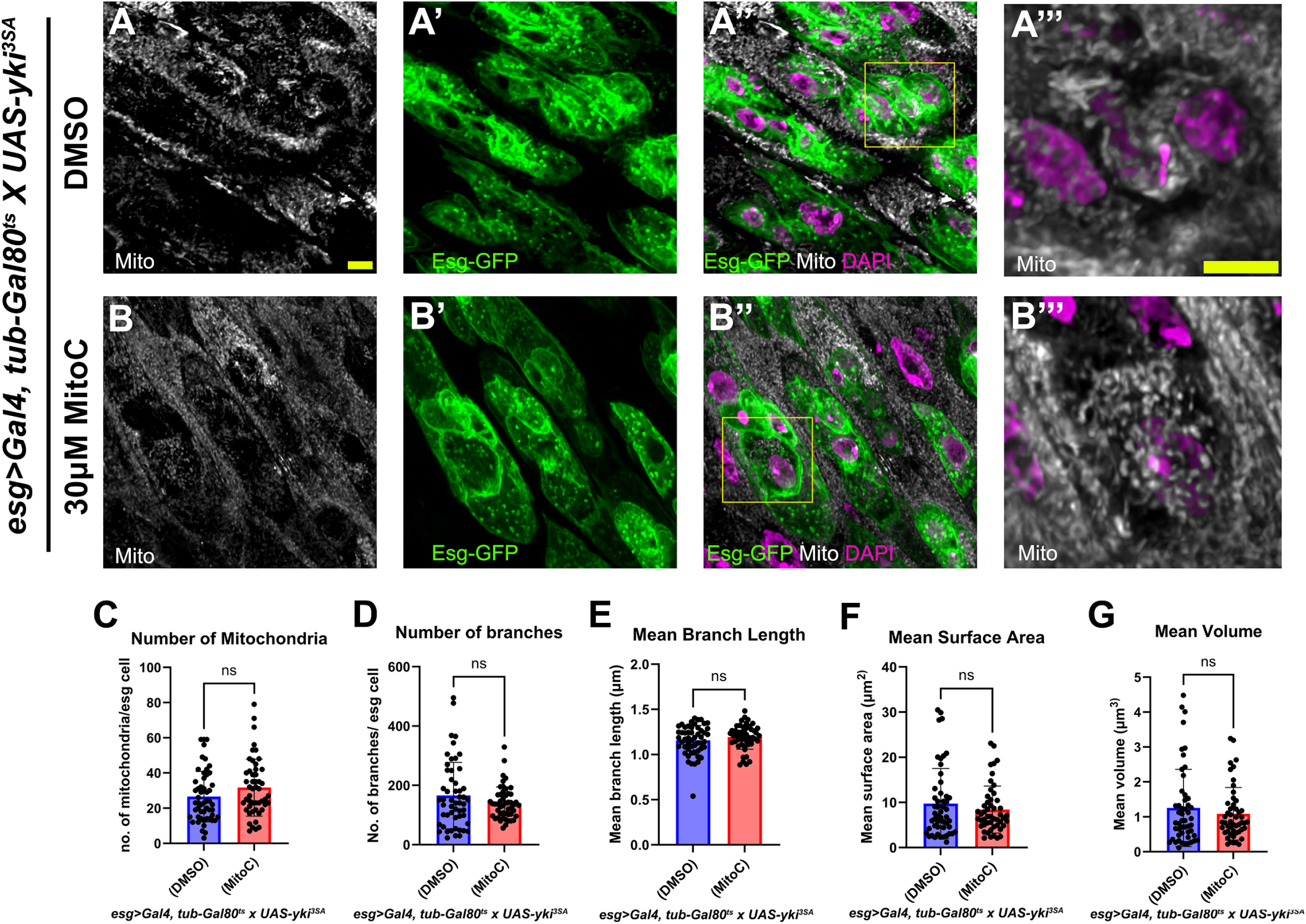
Mitocurcumin does not significantly alter mitochondrial morphology in Yki-driven Esg⁺ tumor cells. Adult midguts from tumor bearing flies stained with MitoView (Gray) upon 30 µM of Mitocurcumin treatment (B, B’, B’’, B’’’) compared to DMSO treatment (A, A’, A’’, A’’’) for 48h. Corresponding graphs of quantitative analysis of mitochondrial morphology parameters: mitochondrial number (C), number of branches (D), mean branch length (E), mean surface area (F), and mean volume (G). Nuclei were stained with DAPI (Magenta in A’’, A’’’, B’’, B’’’). Esg-GFP (Green) marks the intestinal stem cells (ISCs). Data represent mean ± SEM (n = 10 midguts per condition). Statistical analysis was performed using Student’s t-test. p-values indicating statistical significance have been mentioned in the respective graphs. Scale bar: 5 µm (A-B’’’).

### Mitocurcumin disrupts mitochondrial function and shifts tumor metabolic programs

Given the pronounced ROS accumulation observed in mitocurcumin-treated tumors (Fig. 4), we next examined whether redox perturbation was associated with mitochondrial dysfunction. Following 48h of treatment post induction (Fig. 6A), mitochondrial membrane potential was assessed using TMRE staining and flow cytometric analysis of GFP-positive tumor cells. Mitocurcumin-treated samples exhibited a significant reduction in TMRE fluorescence compared to DMSO controls, indicating mitochondrial depolarization (Fig. 6B-C). Total ATP levels in the midguts from mitocurcumin-treated tumor-bearing flies displayed a downward trend as compared to the DMSO control (Fig. 6D). Similarly, mitochondrial abundance, assessed by mtCol expression, showed a modest reduction (Fig. 6E). These findings suggest that mitocurcumin primarily affects mitochondrial function rather than substantially reducing mitochondrial mass and morphology with the 48h treatment window. To further investigate mitochondrial integrity, we analyzed transcriptional changes in components of the electron transport chain (ETC). Mitocurcumin treatment significantly suppressed expression of multiple ETC-associated genes, including Rieske iron-sulfur protein (*RFeSp)*, succinyl-coenzyme A synthetase flavoprotein subunit (*scs-fp)*, Cytochrome c Oxidase subunit (*coxVa)*, and bellwether (*blw)*, whereas mitochondrial acyl carrier protein 1 (*mtACP1)* expression remained unchanged (Fig. 6F–J). The selective downregulation of ETC components is consistent with impaired oxidative phosphorylation and reduced mitochondrial respiratory capacity. In contrast, genes associated with glycolysis and the pentose phosphate pathway (PPP) were upregulated. Expression of Pyruvate kinase (*pyk)* and *pgd* was significantly increased, while *LDH* exhibited a modest upward trend (Fig. 6K–M). Together, these data indicate that mitocurcumin-induced mitochondrial depolarization is accompanied by transcriptional suppression of oxidative phosphorylation and a compensatory shift toward glycolytic and PPP-linked metabolic programs within the Yki-driven tumors.

**Figure 6.**
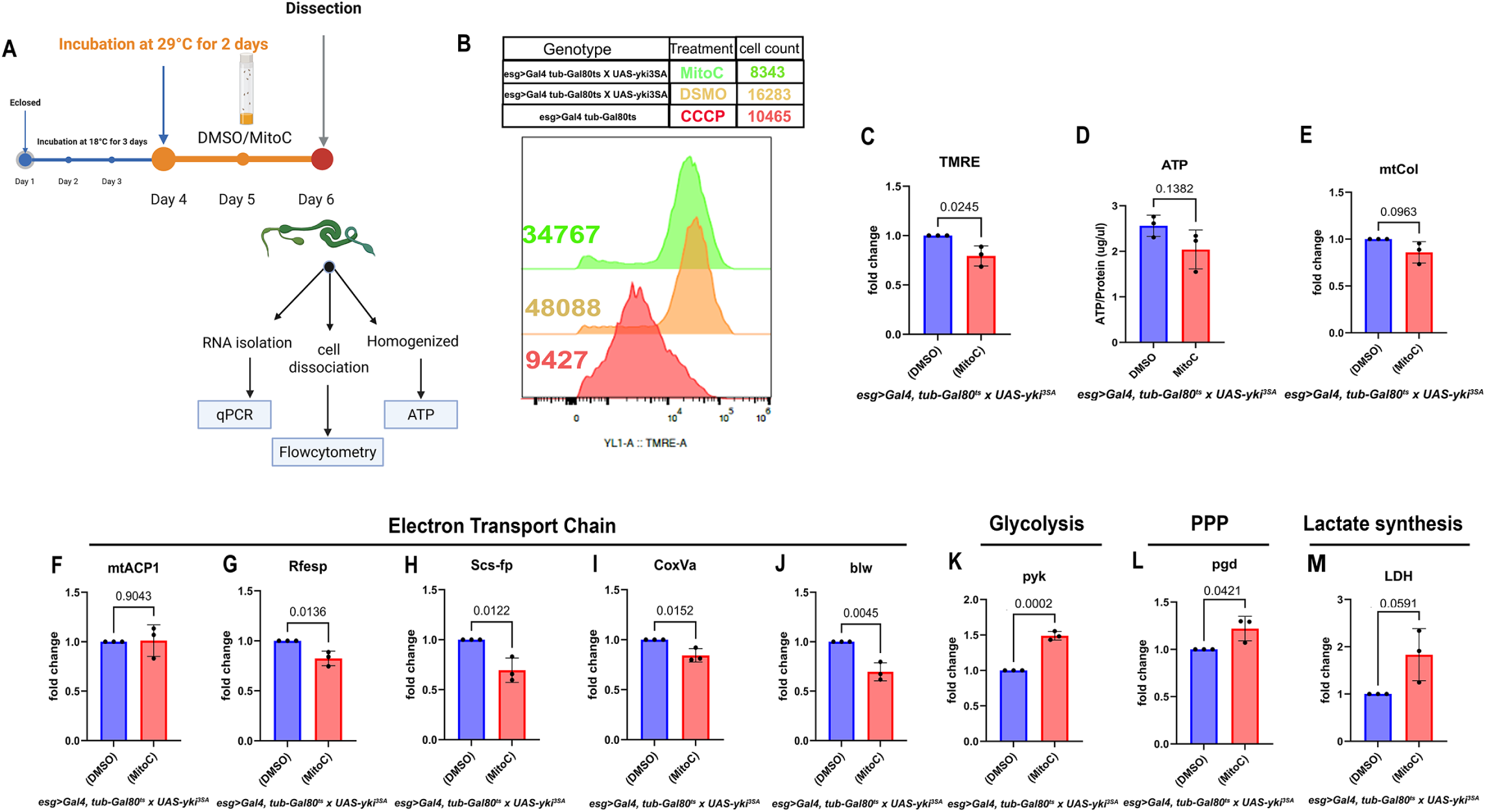
Mitocurcumin disrupts mitochondrial function and shifts tumor metabolic program. Schematic illustration of drug treatment regimen with timeline and experimental setup (A). TMRE-based flow cytometry histograms of GFP-positive tumor cells upon Mitocurcumin treatment compared to DMSO, and CCCP-treated samples. GFP-positive cells were gated for analysis, and the median intensity was indicated respectively (B). Graph showing quantification of TMRE fluorescence intensity (ΔΨm) of Escargot positive cells from the midgut of Mitocurcumin-treated tumor bearing flies compared to DMSO treated tumor bearing flies (C). Graph showing quantification of ATP levels normalized to protein in the midguts upon Mitocurcumin treatment compared to DMSO treatment (D). Graphs showing quantification of the relative mRNA expression of mitochondrial content marker *mt:CoI* (E). Graphs showing quantification of the relative mRNA expression of *mtACP1* (F)*, RFesp* (G)*, scs-fp* (H)*, cox5A* (I)*, blw* (J), *pyk* (K), *pgd* (L), and *ldh* (M) from the midgut of Mitocurcumin-treated tumor-bearing flies compared to DMSO treated tumor bearing flies. RNA was isolated from 10 midguts per replicate (n = 3 biological replicates). Data represent mean ± SEM. Statistical significance was determined using Student’s t-test. p-values indicating statistical significance have been mentioned in the respective graphs. Created in BioRender. Khadilkar, R. (2026) https://BioRender.com/zq9bflj.

### Metabolomic profiling reveals distinct metabolic changes associated with mitocurcumin treatment

Untargeted metabolomic profiling was conducted to examine changes associated with Yki-driven intestinal tumorigenesis and the effects of Mitocurcumin treatment. Peak intensities were log-transformed and scaled to minimize technical variability prior to downstream analysis (Supplementary Fig. S3A). To assess global metabolomic differences between midguts from DMSO-treated Yki tumor bearing flies and Mitocurcumin-treated tumor bearing flies, partial least squares discriminant analysis (PLS-DA) was performed. The resulting score plot showed a clear separation between the two groups, indicating substantial treatment-associated metabolic divergence (Fig. 7A). Hierarchical clustering analysis of significantly altered metabolites further supported this distinction, with samples grouping according to treatment condition (Fig. 7B). Mitocurcumin treatment led to broad metabolic remodelling, affecting multiple metabolite classes, including amino acids, lipid-related species, and intermediates involved in energy metabolism. A total of 51 metabolites were identified as significantly altered between conditions (41 increased and 10 decreased in abundance in Mitocurcumin-treated samples relative to controls), based on a variable importance in projection (VIP) score > 1. Volcano plot analysis further highlighted metabolites exhibiting both statistically significant changes and substantial fold differences (Fig. 7C; Supplementary Fig. S3B). Pathway enrichment analysis of these differential metabolites revealed significant involvement of mitochondrial function, fatty acid metabolism, and amino acid metabolism pathways (Supplementary Fig. S3C–D). To gain deeper insight into mitochondrial alterations, key metabolites associated with energy metabolism were examined (Fig. 7D). Mitocurcumin-treated tumor tissues showed elevated levels of several acylcarnitines, including myristoylcarnitine, palmitoylcarnitine, and acetyl-L-carnitine, alongside increased abundance of tricarboxylic acid (TCA) cycle intermediates such as malate and succinate. These changes are consistent with enhanced mitochondrial metabolic activity following treatment. In addition, increased levels of methionine sulfoxide suggest elevated oxidative stress and disruption of redox balance. Beyond mitochondrial metabolism, alterations were also observed in pathways related to nucleotide turnover and membrane dynamics (Fig. 7E). Midguts from mitocurcumin-treated tumor bearing flies exhibited increased levels of purine metabolites, including inosine, xanthine, and uric acid, indicating enhanced nucleotide metabolism. Conversely, glycerophosphocholine, a metabolite associated with membrane remodelling and cellular proliferation, was reduced following treatment (Fig. 7E). Collectively, these findings indicate that Mitocurcumin influences not only mitochondrial processes but also broader metabolic pathways linked to tumor growth and cellular biosynthesis.

**Figure 7.**
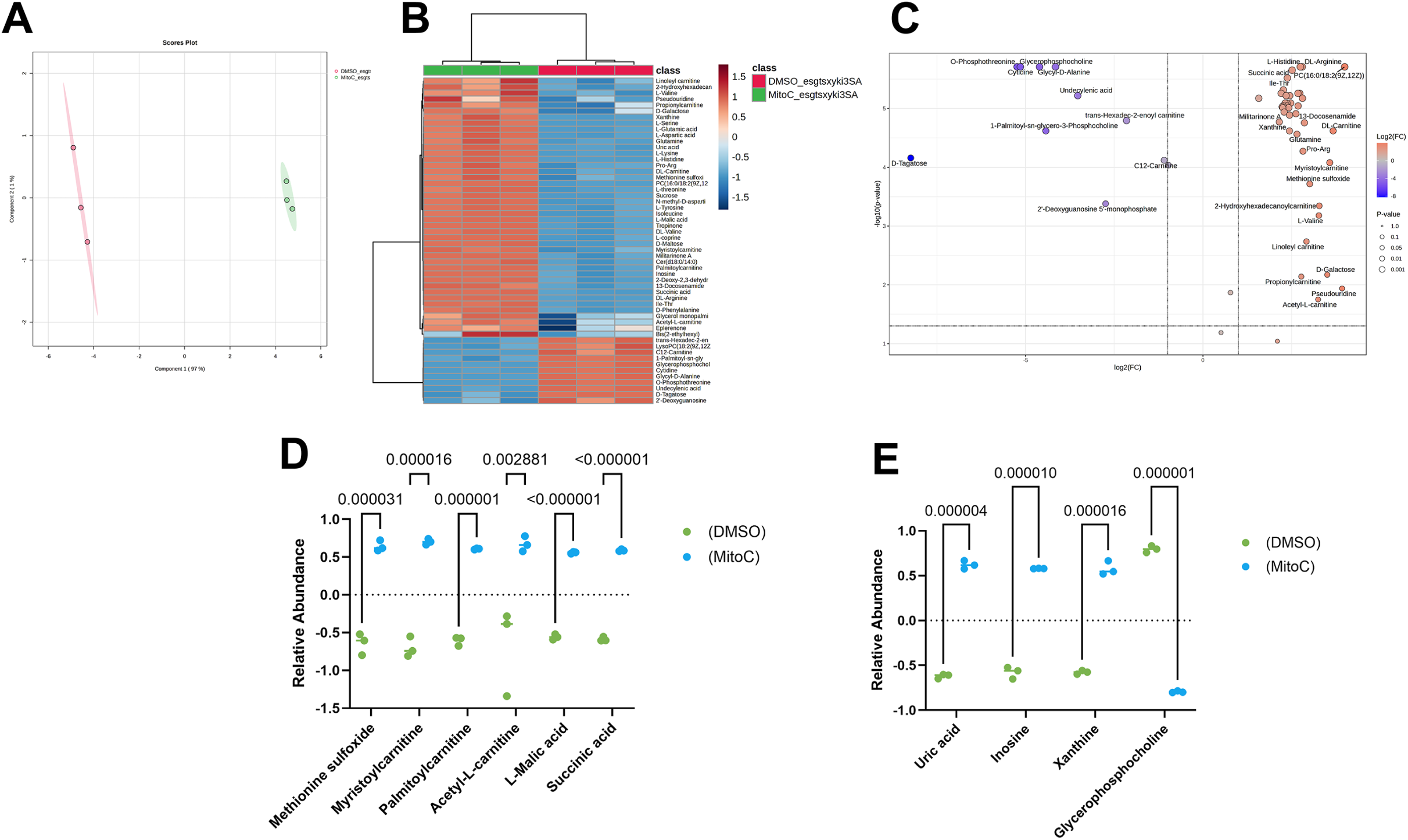
Mitocurcumin induces metabolic reprogramming in Yki-driven gut tumors. Partial least squares discriminant analysis (PLS-DA) score plot of Mitocurcumin- and DMSO-treated samples of tumor bearingflies (A). Heatmap of differentially abundant metabolites with hierarchical clustering (B). Each row represents a metabolite, and each column represents a biological replicate (n = 3).Volcano plot highlighting upregulated and downregulated metabolites (C). Relative abundance of selected metabolites (D, E). Data were log₂-transformed and Pareto-scaled prior to analysis using MetaboAnalyst. Data represent mean ± SEM (n = 3 biological replicates; ∼40 midguts per replicate). Statistical significance was determined using Student’s t-test. p-values indicating statistical significance have been mentioned in the respective graphs.

### Mitocurcumin reduces tumor-associated abdominal bloating and improves overall survival

To assess organism-level consequences of mitocurcumin treatment, gross morphology and lifespan were evaluated in tumor-bearing flies. In order to assess the effect of mitocurcumin at an organismal level we used two different regimens. In the first regimen, we incubated the *esg^ts^-Gal4 X UAS-yki^3SA^* flies at 18°C for 3 days and began induction at 29°C on day 4 (Fig. 8A). Six days post induction we observed a peculiar bloating phenotype in tumor bearing flies treated with DMSO alone. Consistent with previous reports, overexpression of constitutively active Yki resulted in pronounced abdominal bloating in DMSO-treated flies, reflecting severe intestinal dysplasia and compromised gut homeostasis (Song et al., 2019) (Fig. 8B). In contrast, mitocurcumin-treated flies exhibited a marked reduction in abdominal distension (Fig. 8C). Quantitative analysis revealed a significant decrease in the proportion of bloated individuals following mitocurcumin treatment (Fig. 8D). In the second regimen, we incubated the *esg^ts^-Gal4 X UAS-yki^3SA^* flies at 18°C for 3 days and began induction at 29°C on day 4. We began treatment with mitocurcumin and DMSO control six days post-induction when bloating was observed in all the tumor bearing flies for 48h to assess if there is any visible rescue and reversibility of the abdominal bloating (Fig. 8E). Gross morphological examination upon 48h treatment with mitocurcumin or DMSO confirmed visible improvement in the bloating phenotype abdominal architecture upon treatment (Fig. 8F-G). Measurement of the abdomen-to-head (A/H) area ratio demonstrated a substantial reduction in abdominal enlargement in mitocurcumin-treated flies compared with DMSO controls (Fig. 8H). Finally, survival analysis showed that mitocurcumin significantly extended lifespan in tumor-bearing flies relative to DMSO-treated controls, increasing median survival from 28 days to 36 days (Fig. 8I). Together, these findings demonstrate that mitocurcumin-mediated tumor regression translates into improved intestinal physiology and enhanced organismal survival, indicating that redox-targeted intervention alleviates both local tumor burden and systemic consequences of Yki-driven hyperplasia in the midgut (Fig. 9).

**Figure 8.**
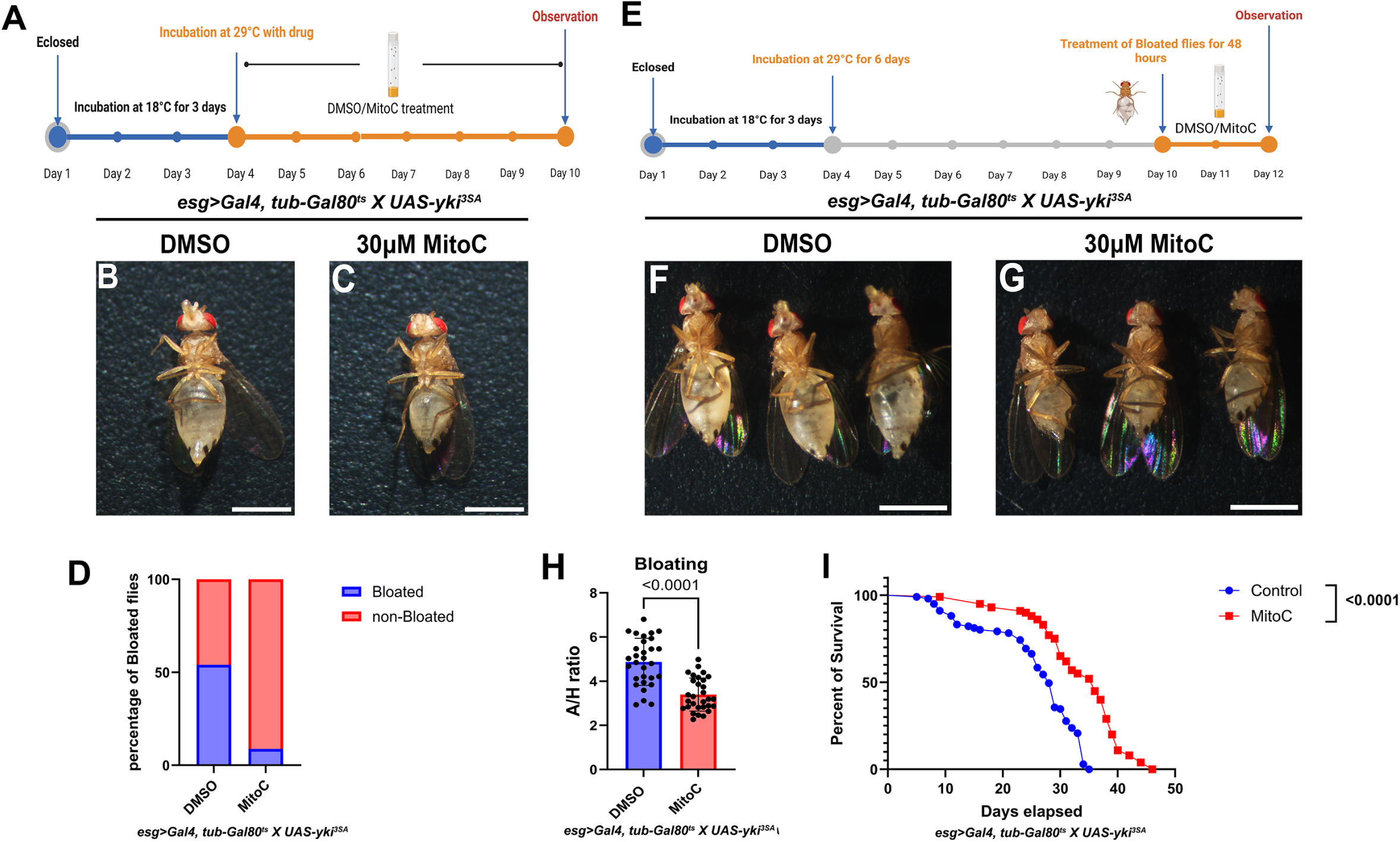
Mitocurcumin reduces tumor-associated abdominal bloating and improves overall survival. Schematic illustration of drug treatment regimen with simultaneous induction of the *yki^3SA^* transgene from Day 4 for phenotypic assessment (A). Representative images of tumor-bearing adult flies upon Mitocurcumin (MitoC) treatment compared to DMSO treatment with this experimental regimen(B-C). Quantification of the proportion of bloated flies with this experimental regimen (D). Schematic illustration of drug treatment regimen for 48h post six days of induction of the *yki^3SA^* transgene from Day 4 for phenotypic assessment (E). Representative images of tumor-bearing adult flies upon Mitocurcumin (MitoC) treatment compared to DMSO treatment with this experimental regimen (F-G). Quantification of abdominal bloating expressed as abdomen-to-head (A/H) area ratio (H). Kaplan–Meier survival curves of tumor-bearing flies treated with Mitocurcumin (30 µM) compared to DMSO treatment (I). Survival analysis was performed using 100 flies per condition (4 independent vials containing 25 flies each). Data represent mean ± SEM. Statistical significance was determined using Student’s t-test (F) and the log-rank (Mantel–Cox) test (G). Scale bar: 1 mm (A-E). p-values indicating statistical significance have been mentioned in the respective graphs. Created in BioRender. Khadilkar, R. (2026) https://BioRender.com/0drozcc. Created in BioRender. Khadilkar, R. (2026) https://BioRender.com/kd8xwi3

**Figure 9.**
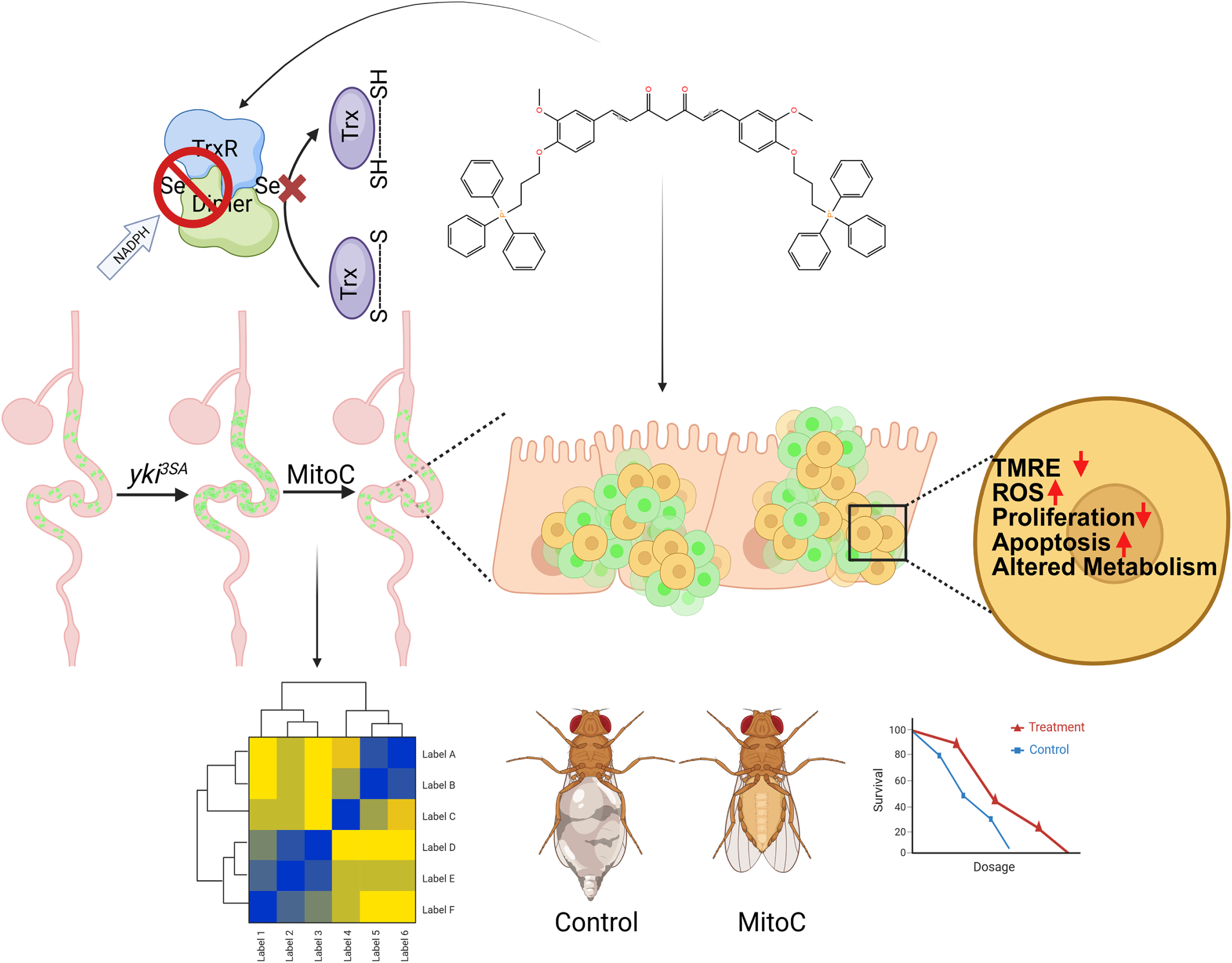
Graphical Summary. Schematic model illustrating Mitocurcumin-mediated inhibition of thioredoxin reductase, leading to increased oxidative stress, metabolic reprogramming, and reduction in tumor burden of Yki-driven intestinal tumors. Created in BioRender. Khadilkar, R. (2026) https://BioRender.com/zzjjg3g

## Discussion

Dysregulated redox homeostasis is a crucial vulnerability of cancer cells, and recent reports have shown how to exploit this vulnerability to selectively kill them (Li et al., 2023; Park et al., 2026; Xing et al., 2022). Cancer cells typically operate under high oxidative stress, which supports tumor growth and treatment resistance. Paradoxically, excessive accumulation of ROS leads to cell death; hence, redox-based therapeutic strategies are emerging(Chen et al., 2025). Mitocurcumin, a triphenylphosphonium-conjugated derivative of curcumin, was previously shown to accumulate in mitochondria and to kill lung cancer cells largely through inhibition of mitochondrial thioredoxin reductase 2 (TrxR2) and consequent ROS-driven apoptosis(Jayakumar et al., 2017). Our work provides an in vivo validation using a genetically tractable *Drosophila* intestinal tumor model, in which we observe a potent accumulation of ROS following Mitocurcumin treatment, resulting in increased apoptosis of tumor cells and a consequent decrease in tumor burden. In the *Drosophila* midgut, ROS acts as a vital signaling rheostat; moderate levels promote intestinal stem cell (ISC) proliferation, while excessive levels trigger apoptosis (Morris & Jasper, 2021; Mundorf et al., 2019; Zhang et al., 2024). Our study demonstrates that mitocurcumin not only results in dysregulated redox homeostasis and mitochondrial metabolism but also ameliorates the typical abdominal bloating phenotype reported earlier in *Drosophila* tumor models and also improves the survival of the tumor-bearing flies upon treatment (Ding et al., 2021; Kwon et al., 2015; Song et al., 2019).

Strikingly, our comparative docking, molecular dynamics, and MM-GBSA analyses show a robust in silico binding of Mitocurcumin to *Drosophila* TrxR1. Mammalian TrxR relies on a C-terminal selenocysteine (Sec) residue for its catalytic activity, which is the canonical target for many electrophilic inhibitors(Jayakumar et al., 2017). In contrast, *Drosophila* TrxR utilizes a cysteine (Cys) residue while retaining high catalytic efficiency(Missirlis et al., 2002). Our data suggests Mitocurcumin’s inhibitory potential does not strictly require the presence of a Sec residue. Instead, the drug effectively anchors into a conserved structural pocket near the FAD-binding site, stabilized by π-stacking with Tyr197. While direct in vitro enzymatic kinetics for *Drosophila* TrxR1 have yet to be characterized, our 500 ns MD simulations, coupled with the profound intracellular ROS surge, provide compelling in vivo evidence of functional impairment of TrxR. We show that mitocurcumin reduces Yki-induced intestinal tumors in a dose-dependent manner, with a clear reduction in mitotically active cells and a corresponding increase in apoptotic cDcp1-positive cells within Esg-positive cell clusters. This demonstrates that the anti-proliferative and pro-apoptotic activity of mitocurcumin observed in vitro is reproducible in an intact, oncogene-driven tumor in vivo. We observe that increased ROS triggers the anti-oxidant response pathway by upregulating Cnc (the *Drosophila* Nrf2 homolog) and downregulating Keap1. However, this activation does not result in a robust upregulation of downstream antioxidant enzyme transcripts. It is possible that although transcript levels are not affected, the activity of antioxidant enzymes and other ROS-quenching machinery could be altered by mitocurcumin treatment, which warrants further investigation. The mechanism underlying the compensatory antioxidant machinery could be further explored to elucidate the intricacies of this fine balance in future studies following mitocurcumin treatment. We also showed that the mitochondrial network morphology of these Esg-positive cell clusters was comparable across treatment groups, even though TMRE staining showed clear mitochondrial depolarization and the transcript levels of ETC complex genes were suppressed. It could mean the functional impairment of mitochondria genuinely precedes any structural remodelling of the mitochondria within this drug treatment time window. Also, there is a possibility that the most severely damaged mitochondria are already turned over during this time window of drug treatment and hence what we measure is a surviving population of mitochondria whose energy metabolism is completely rewired.

Mitocurcumin treatment reprograms tumor metabolism by selectively suppressing components of the electron transport chain, consistent with impaired oxidative phosphorylation. The metabolism shifts towards increased glycolysis and upregulation of genes related to the pentose phosphate pathway. Now, stress conditions that drive increased glycolysis can rewire mitochondrial metabolism by leading to succinate accumulation, which promotes reverse electron transfer and thereby promotes ROS(Erlich et al., 2022). Our metabolomics analysis shows a clear succinate accumulation upon mitocurcumin treatment. Perturbation of mitochondrial function could increase PPP activity and elevate mitochondrial superoxide and intracellular hydrogen peroxide, linking the PPP to a higher ROS state(Hambardikar et al., 2022). Now, in activated macrophages and other immune cells, it has been shown that an increased glycolysis-PPP shunt specifically supports increased ROS/superoxide/nitric oxide output by elevating NADPH availability(Patra & Hay, 2014; Teslaa et al., 2023). Untargeted metabolomic profiling reinforced this picture: PLS-DA and hierarchical clustering showed clear separation between DMSO- and mitocurcumin-treated tumors, with 51 metabolites significantly altered, with 41 upregulated and 10 downregulated metabolites.. Treated tumors accumulated acylcarnitines (myristoylcarnitine, palmitoylcarnitine andacetyl-L-carnitine) and TCA-cycle intermediates (malate, succinate), alongside elevated methionine sulfoxide indicative of oxidative stress, while purine metabolites (inosine, xanthine, uric acid) rose and the proliferation-associated lipid glycerophosphocholine declined — together giving a clear indication of disrupted mitochondrial energy metabolism coupled with altered nucleotide turnover and membrane remodelling. These molecular and metabolic perturbations translated into meaningful organism-level benefits: mitocurcumin markedly reduced the tumor-associated abdominal bloating characteristic of Yki-driven gut dysplasia(Ding et al., 2021; Kwon et al., 2015; Song et al., 2019), partially reversed bloating even when treatment began after bloating had already set in, and significantly extended the survival of tumor-bearing flies, raising the median lifespan from 28 to 36 days. Collectively, these results indicate that mitocurcumin’s redox- and metabolism-targeted mechanism of action is not confined to the cellular level but produces tangible improvements in gut physiology and whole-organism health in a Yki-driven tumor model. The systemic signalling and mechanism underlying the rescue of abdominal bloating and cachectic phenotypes needs detailed exploration. The mechanistic role of Mitocurcumin in facilitating these systemic changes, particularly in cachexia models, can be explored further to understand the inter-organ signaling driven by altered metabolism.

Overall, our study provides evidence for mitocurcumin as a therapeutic candidate that targets the tumor burden by altering ROS and metabolism in the tumor cells and opens interesting future directions for understanding how targeting mitochondria in tumor cells can have organism-wide systemic effects on the tumor macro-environment.

## Acknowledgements

We would like to thank the Digital Imaging Facility (DIF) and the Common Instrumentation Facilities (CIF) at ACTREC for all the support. We thank the Bloomington Drosophila Stock Center, Developmental Studies Hybridoma Bank, and the fly community for the fly stocks. We are thankful to the Stem Cell and Tissue Homeostasis lab for useful input and discussions.

## Author contributions

Conceptualization: R.J.K.; Data curation: R.J.K., S.D., U.C., S.J.; Formal analysis: S.D., U.C., N.G.; Funding acquisition: R.J.K.; Investigation: S.D., U.C., N.G., S.J.; Methodology: S.D., U.C., N.G., S.J.; Project administration: R.J.K.; Resources: R.J.K. V.G.; Supervision: R.J.K.; Validation: S.D., U.C.; Visualization: S.D., U.C., R.J.K.; Writing – original draft: S.D., U.C., R.J.K.; Writing – review & editing: S.D., U.C., R.J.K.

## Funding

This study was funded by Department of Biotechnology, Ministry of Science and Technology, India for the Har Gobind Khorana – Innovative Young Biotechnologist Award (no. BT/13/IYBA/2020/14) to R.J.K., Ramalingaswami Re-entry Fellowship from the Department of Biotechnology, Ministry of Science and Technology, India (BT/RLF/Re-entry/19/2020) to R.J.K. This work was also funded by a Basic and Translational Research in Cancer grant (no.1/3(7)/2020/TMC/R&D-II/8823 Dt. 30.07.2021), Capacity Building and Development of Novel and Cutting-edge Research Activities (no.1/3(4)/2021/TMC/R&D-II/15063 Dt. 15.12.2021) from the Department of Atomic Energy, Government of India. The LC-MS system was procured through the CAR grant by Indian Council of Medical Research (ICMR/BMS/Phase-1/CAR/2023) awarded to V.G.

## Data availability

All relevant data can be found within the article and its supplementary information

## Competing interests

The authors declare no competing or financial interests.

## Materials and Methods

### Fly stocks

*w^1118^*, UAS-*yki^3SA^* (BL28817, RRID: BDSC_28817), and *esg-Gal4, UAS-GFP; tub-Gal80^ts^*

### *Drosophila* Husbandry and tumor induction

All fly stocks and crosses were maintained on standard cornmeal agar medium under a 12 h light/12 h dark cycle. To suppress Gal4 activity during development, progeny were maintained at 18 °C until eclosion and further incubated at 18°C for 3 days to allow normal midgut maturation post eclosion. Subsequently, flies were shifted to 29°C to induce *yki^3SA^* expression specifically in esg-positive intestinal stem cells, as previously described (Leung et al., 2024). The transgene expression for tumor induction at 29°C was followed as per experimental regimen described in respective figures. For all experimental analysis, virgin female adult flies were used to ensure consistency and minimize variability.

### Sequence and Structural Analysis of Thioredoxin Reductase

Protein sequences of thioredoxin reductase (TrxR) from *Drosophila melanogaster* (TrxR1; P91938), *Rattus norvegicus* (TrxR2; Q9Z0J6), and *Homo sapiens* (TrxR1; Q16881) were retrieved from the UniProt database. Multiple sequence alignment was performed using Clustal Omega, and the resulting alignment was visualized and analyzed using Jalview to assess conservation of functional domains, including the catalytic core and cofactor-binding regions.

### Thioredoxin Reductase and Mitocurcumin preparation using Schrödinger

The three-dimensional structures of TrxR homologs were obtained from the Protein Data Bank. The crystal structure of *Drosophila melanogaster* TrxR1 (PDB ID: 3DH9) and the structure of mammalian TrxR (PDB ID: 3QFA) were superimposed using the Maestro tool (RRID:SCR_016748) of Schrödinger (RRID:SCR_014879) for the comparison of overall fold and active-site architecture. The 3D structures of Thioredoxin Reductase were prepared by assigning protonation state at pH (7.4 ± 0.5), adding hydrogen bonds and minimising structures using OPLS4 force field. The 2D structure of curcumin was retrieved from the PubChem database (PubChem CID: 969516). The 2D structure of Mitocurcumin (MitoC) was sketched by adding lipophilic triphenylphosphonium group to the curcumin using Maestro and subjected to energy minimization. Both curcumin and Mitocurcumin were prepared using the LigPrep tool from the Schrödinger suite to generate low-energy three-dimensional conformations by optimising ionisation states at physiological pH (7.4 ± 0.5).

### Molecular Docking of curcumin and Mitocurcumin with Thioredoxin Reductase

Molecular docking was performed using the Glide (RRID:SCR_000187) module within the Schrödinger software suite. Prepared ligands were docked into the predicted active-site pocket of the respective TrxR structures: *Drosophila melanogaster* TrxR1 (PDB ID: 3DH9) and human TrxR1 (PDB ID: 3QFA). Docking grids were generated around the catalytic site, specifically centered on the FAD cofactor within the conserved FAD/NADPH-binding region to encompass the entire active site pocket. Docking calculations were carried out using the standard precision (SP) mode and Glide scores were used to evaluate and compare predicted binding affinities of ligands across TrxR homologs.

### Molecular Dynamics Simulation of the Thioredoxin Reductase – Mitocurcumin docked complexes

Molecular dynamics (MD) simulations were performed to evaluate the stability of the docked Mitocurcumin–TrxR complexes using the Desmond simulation package implemented in Schrödinger. The complexes were embedded in an explicit solvent environment using the TIP3P water model and neutralized by the addition of appropriate counterions. Simulations were conducted under periodic boundary conditions using the OPLS4 force field. Following energy minimization and equilibration, production simulations were carried out for 500ns under constant temperature and pressure conditions. Trajectories were analyzed for backbone RMSD, ligand RMSD, residue-wise fluctuations (RMSF), and protein–ligand contact stability. The binding free energy estimation between the Mitocurcumin-TrxR complexes were performed using the Molecular Mechanics/Generalized Born Surface Area (MM/GBSA) method using ‘thermal_mmgbsa.py’ script from Schrodinger. Representative frames extracted from the MD trajectories were used to calculate the average binding free energy (ΔG), providing an estimate of the energetic favourability of Mitocurcumin binding to TrxR.

### Mitocurcumin Treatment

A 10 mM stock solution of Mitocurcumin (MitoC) was prepared in dimethyl sulfoxide (DMSO; Sigma-Aldrich, D8418). The stock solution was diluted in standard cornmeal agar food to obtain final concentrations ranging from 10µM to 40µM. Control food contained an equivalent volume of DMSO. Drug-containing food was mixed thoroughly, allowed to solidify, and provided to flies. Based on dose–response analyses, 30µM Mitocurcumin was used for all subsequent experiments unless otherwise specified.

### Regimen for Tumor Induction and Drug Treatment

For tumor induction, *esg-GAL4, UAS-GFP; tub-GAL80^ts^ > UAS-yki^3SA^* flies were maintained at 18 °C for 3 days post-eclosion to suppress GAL4 activity and to allow midgut maturation. Flies were then shifted to 29 °C to induce Yki-driven tumor formation in Esg-positive intestinal stem cells. Flies were transferred to food containing either DMSO or mitocurcumin and maintained at 29 °C for 48h or longer or were treated with DMSO or mitocurcumin a few days post tumor induction depending on the experimental regimen mentioned for each experiment in respective figures.

### Regimen for Bloating and Phenotypic Analysis

For bloating analysis, two experimental regimens were employed:

Concurrent induction and treatment: Flies were maintained at 29 °C for 6 days in the presence of DMSO or Mitocurcumin to assess tumor-associated abdominal bloating.

Post-induction treatment: Flies were first subjected to tumor induction at 29 °C for 6 days to establish the bloating phenotype, followed by transfer to DMSO- or Mitocurcumin-containing food for an additional 48h to evaluate therapeutic reversal of bloating.

### Immunofluorescence and Tissue Staining

Adult midguts were dissected in cold phosphate-buffered saline (PBS) and fixed in 4% paraformaldehyde (PFA) for 30 minutes at 29 °C. Tissues were permeabilized and blocked in PBST (0.1% Triton X-100 in PBS) and 20% normal goat serum. Primary antibody incubation was performed overnight at 4 °C using antibodies against phospho-Histone H3, pH3 (Sigma-Millipore, 06-570) to label proliferating cells and cleaved *Drosophila* Dcp1, cDCP1 (Asp216) to detect apoptotic cells. After washing, tissues were incubated with appropriate fluorescent secondary antibodies and counterstained with DAPI to visualize nuclei. Alexa-Fluor 568 conjugated secondary antibodies – Goat-raised anti-rabbit 568 (1:400, Invitrogen, RRID: AB_143157) were used for immunofluorescence-based experiments. The midguts were mounted in VECTASHIELD mounting medium (Vector Laboratories; RRID: AB_2336790).

### Confocal Microscopy and Image Analysis

Images were acquired on Zeiss LSM 980, Leica SP8 or Nikon AX confocal microscopes. The raw image files were processed in ImageJ/Fiji software (v1.54d, RRID: SCR_003070) and converted to TIFF format (RGB mode), with individual or merged channels generated as required. TIFF files were subsequently imported into Adobe Photoshop (RRID: SCR_014199) and assembled into figure panels on a RGB canvas. Final figures were saved in TIFF format using LZW compression at a resolution of 600 dpi or more.

### Tumor Burden Quantification

Whole-gut tile-scan Z-stack images were acquired on Leica SP8 confocal microscope (20X objective) using 512×512 frame size per tile with no averaging and a z-step size of 1.5 microns and identical acquisition settings across conditions. The images were stitched using an in-built application of Leica Application Suite X (RRID:SCR_013673). Three-dimensional reconstruction and volumetric analysis were performed using Imaris software (RRID:SCR_007370). esg-positive (GFP-positive) tumor volume was calculated using the *Surface* creation module and normalized to total gut volume, delineated by DAPI staining. Tumor burden was expressed as the ratio of esg-positive volume to total gut volume.

### cDCP1 Quantification in esg-Positive Tumor Cells

The images were acquired on Zeiss LSM 980 using Airyscan mode (40x objective, 1 zoom, oil immersion). Quantitative analysis of apoptosis was performed using Imaris software (RRID:SCR_007370). esg-positive tumor cells were segmented using the *Surface* creation module based on GFP fluorescence. cDCP1-positive apoptotic puncta were detected using the *Spots* module with identical detection parameters applied across all samples.

To specifically quantify apoptosis within tumor cells, only cDCP1 puncta localized within esg-positive surfaces were included in the analysis. Apoptotic burden was calculated as the number of cDCP1 puncta normalized to total esg-positive area, and expressed as cDCP1 puncta per esg-positive area. This approach enabled cell-type–specific quantification of apoptosis within the tumor compartment.

### Reactive Oxygen Species (ROS) Measurement

Intracellular ROS levels were measured using CellROX™ Deep Red (Thermofisher scientific, C10422) following the manufacturer’s protocol. Briefly, the midguts were dissected in Schneider’s insect medium (Thermofisher scientific, 21720024). Dissected midguts were incubated with 5µM CellROX reagent at 29 °C for 30 min, followed by fixation in 4% PFA. 15 mM Paraquat (PQ) (Sigma 36541) treatment was used as a positive control for ROS induction. The images were acquired on Zeiss LSM 980 using confocal mode (40x objective, 0.6 zoom, oil immersion). Fluorescence intensity was quantified specifically within esg-positive tumor clusters using ImageJ/FIJI, and expressed as corrected total cell fluorescence (CTCF).

### Gene Expression Analysis by Quantitative RT-PCR

Total RNA was extracted from 10 dissected midguts per condition using the TRIzol reagent. Briefly, the dissected guts were lysed in TRIzol (Thermo Fisher Scientific, 11596018). RNA was isolated from the aqueous layer post-chloroform treatment, followed by RNA isolation according to the manufacturer’s protocol. RNA yield was quantified using Nanodrop. 1µg of mRNA was reverse transcribed using oligo-dT primers (Promega, C110A) and ImProm-II (Promega, A3800). Quantitation of the mRNA transcripts was done using SYBR green chemistry (Thermo Fisher Scientific, Cat. no.: 4367659) in the Quantstudio 5 RT PCR system (Thermo Fisher Scientific) in quadruplets of 10µl reaction. The data were analyzed using the ΔΔCt method and relative mRNA expression was normalized to rp49. Fold change calculations were done in comparison to the DMSO control group. The experiment was done in biological triplicates and statistical analysis was performed using one-way ANOVA (Dunnett) for comparison of all test genotypes with the wild-type control genotype.

## List of primers

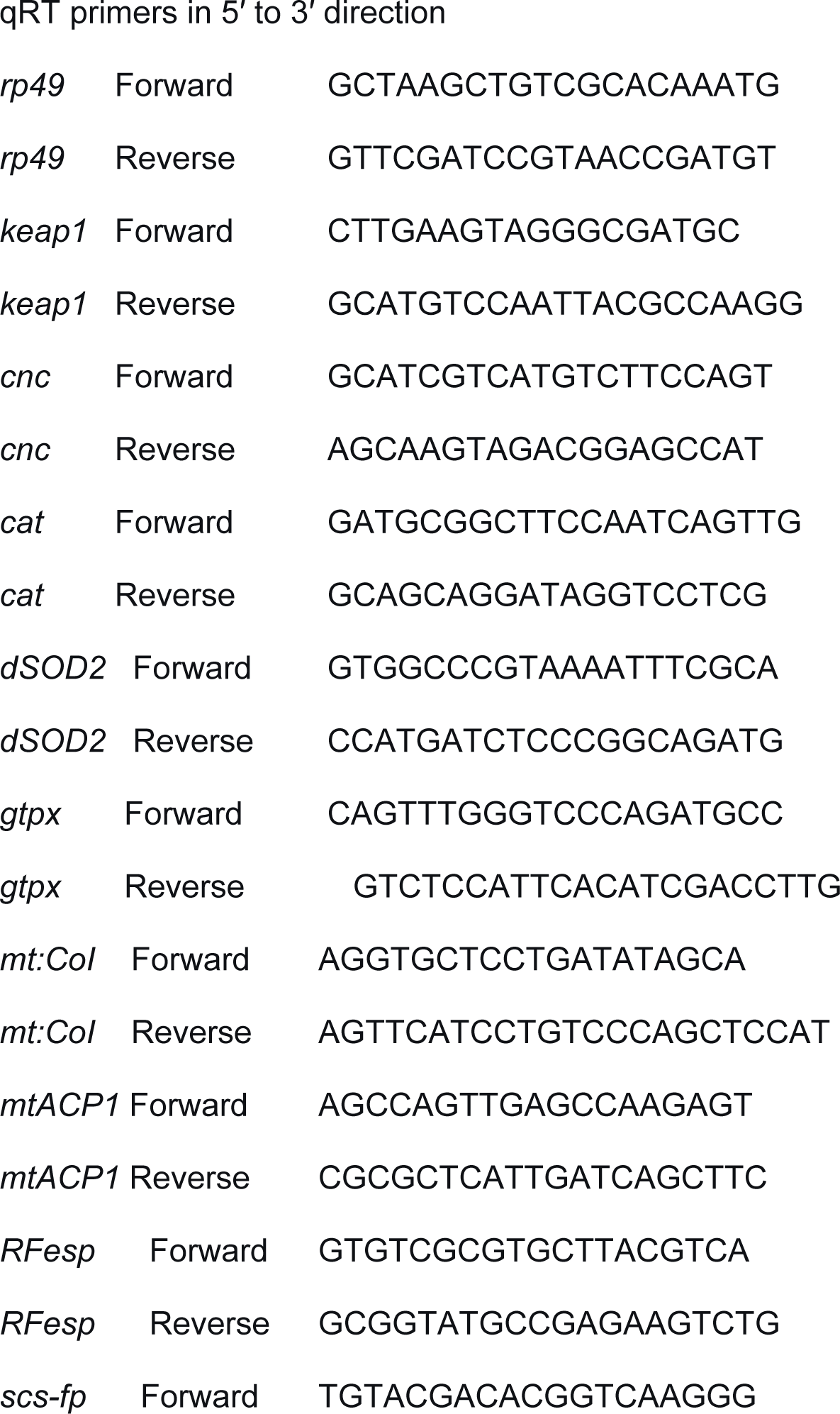
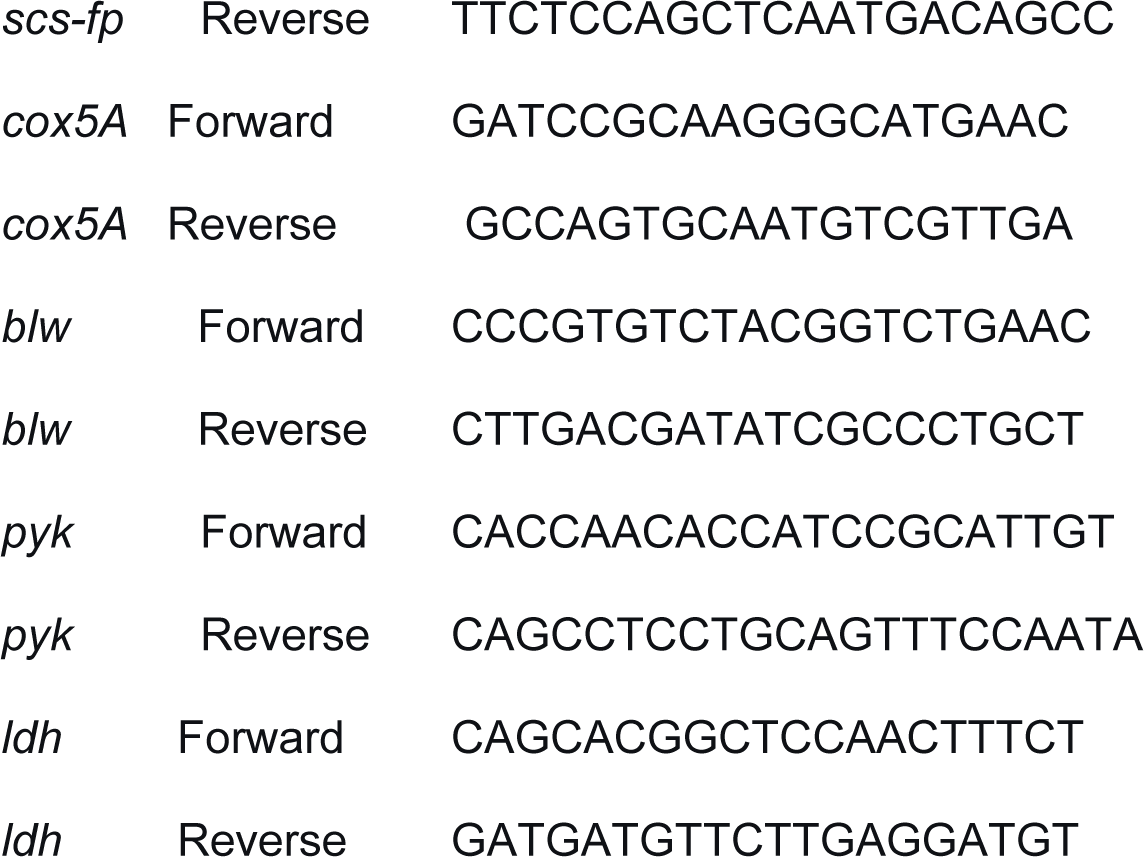

### Mitochondrial Membrane Potential (ΔΨm) Assay

Mitochondrial membrane potential was assessed using TMRE (Thermo Fisher Scientific, T669) staining followed by flow cytometry as described in (Zhang et al., 2022). Dissected guts were dissociated into single-cell suspensions using Collagenase type I (1:1000 of 100U/ml stock solution, Gibco, 17100017) and EDTA (final concentration of 2mM) and samples were incubated for 1 hour at 29°C. Samples were vigorously pipetted 30 times using a cut tip every 15 mins to help dissociation. After dissociation, samples were centrifuged at 600g for 10 mins. Supernatant was discarded and 500µl of cold fresh nuclease-free PBS was added. Cells were re-suspended and filtered by 40µm cell strainer (Flowmi, 136800040). The dissociated cells were incubated with 200nM TMRE. Carbonyl cyanide m-chlorophenyl hydrazone, CCCP (Sigma, C2759-100MG) was used as a positive control for mitochondrial depolarization. *w1118* cells and unstained samples served as additional controls. Attune flow cytometer (Thermo Fisher Scientific) was used to acquire the signal. GFP-positive tumor cells were gated for analysis. FlowJo analyzer (RRID:SCR_008520) was used to quantitate signal.

### ATP Measurement

Total ATP levels were quantified from dissected gut tissues using a luminescence-based ATP assay kit (Molecular Probes ATP kit; A22066). As described in (Tennessen et al., 2014) with slight modifications, gut tissues were dissected from 10 tumor bearing flies per biological replicate for both DMSO-treated controls and Mitocurcumin (MitoC) treated samples. Samples were homogenized on ice in 100µL of chaotropic homogenization buffer (6 M guanidine HCl, 100mM Tris-HCl pH 7.8, 4mM EDTA) using a pellet pestle. To normalize ATP levels, 10µL of the homogenate was reserved for protein quantification using a Pierce BCA assay (Catalog number: 23225). The remaining homogenate was boiled for 5 minutes to inactivate ATPases, followed by centrifugation at maximum speed (∼20,000 × g) for 3 minutes at 4°C. 10µL of the supernatant was transferred to wells of a white opaque 96-well plate. The luciferase reaction mix (prepared fresh with reaction buffer, D-luciferin, DTT, and firefly luciferase according to the manufacturer’s instructions) was added (100µL per well) using a multichannel pipette. Luminescence was immediately measured using a plate reader. Three sequential readings were acquired for each well and averaged.

### Mitochondrial Morphology Analysis

Dissected midguts were incubated in Schneider’s insect medium (Thermofisher scientific, 21720024) containing MitoView fix 640 dye for 30 minutes at 29 °C, followed by fixation in 4% PFA. The images were acquired on Nikon AX microscope (100x objective, 3 zoom, oil immersion). Confocal images were analyzed using the Mitochondria Analyzer (RRID:SCR_027707) plugin (https://github.com/AhsenChaudhry/Mitochondria-Analyzer) in FIJI/ImageJ to quantify mitochondrial number, number of branches, mean branch length, mean surface area, and mean volume (Chaudhry et al., 2020).

### Sample Preparation and Metabolite Extraction

For metabolomic profiling, *esg-GAL4, UAS-GFP; tub-GAL80^ts^ > UAS-yki^3SA^* flies were maintained at 29 °C for 6 days in the presence of either DMSO or Mitocurcumin. Following treatment, midgut tissues were dissected and processed for metabolite extraction and LC– MS/MS analysis.

Adult *Drosophila* midgut tissues were obtained from e*sg-GAL4, UAS-GFP; tub-GAL80ts > UAS-yki^3SA^* flies under two experimental conditions: DMSO-treated controls and Mitocurcumin-treated groups. For each biological replicate, approximately 40 midguts were carefully dissected in chilled phosphate-buffered saline (PBS) to limit metabolic alterations during tissue collection. A total of three independent biological replicates were prepared per condition. Immediately after dissection, tissues were transferred into pre-cooled ethanol to rapidly halt ongoing metabolic activity. All subsequent handling steps were conducted at low temperature to preserve metabolite stability. Tissue disruption was carried out using a bead-based homogenization approach. Samples were placed in screw-cap tubes containing zirconium beads and processed using a bead beater under cold conditions to ensure efficient cellular lysis.

Following homogenization, samples were centrifuged to separate insoluble material, and the metabolite-containing supernatant was collected. Extraction was performed using a cold methanol:water mixture (4:1, v/v), with volumes adjusted proportionally to tissue input. To maximize metabolite recovery, extracts were incubated at −20 °C for 1 hour prior to centrifugation at 16,000 × g for 15 minutes at 4°C. The clarified supernatant was transferred to fresh tubes and evaporated using a vacuum concentrator. Dried residues were subsequently reconstituted in methanol:water (1:1, v/v) and normalized to the initial tissue amount. Reconstituted samples were briefly vortexed for 5 minutes and centrifuged again under the same conditions to remove any remaining particulates. The final supernatant was transferred into LC-MS vials for downstream analysis.

### LC–MS/MS Analysis

Metabolomic profiling was conducted using an ultra-performance liquid chromatography (UPLC) system coupled to a high-resolution Orbitrap mass spectrometer. Polar metabolites were separated using a hydrophilic interaction liquid chromatography (HILIC) column. Data acquisition was performed in both positive and negative electrospray ionization modes to broaden metabolite coverage. Mass spectra were collected over an m/z range of 50–1000 Da, incorporating both full-scan MS and data-dependent MS/MS analyses. Scan times were set at 500 ms for MS and 130 ms for MS/MS events, with collision energies ranging between 10 and 50 eV. All solvents utilized were of LC-MS grade to maintain analytical consistency.

### Data Processing, Metabolite Identification, and Statistical Analysis

Raw LC-MS datasets were processed using Compound Discoverer (version 3.4, Thermo Fisher Scientific). The workflow included feature detection, retention time correction, alignment across samples, and compound annotation through comparison with spectral libraries and databases.

Processed data were further analyzed using MetaboAnalyst (https://www.metaboanalyst.ca/, RRID:SCR_015539). To improve comparability and reduce technical variability, datasets were subjected to log transformation followed by scaling. Multivariate statistical methods, including principal component analysis (PCA), were applied to evaluate global metabolic differences between experimental groups. Hierarchical clustering was used to examine patterns in metabolite abundance across samples.

Differential metabolites were identified based on fold-change thresholds and statistical significance criteria. Visualization techniques, including heatmaps and volcano plots, were employed to illustrate metabolomic alterations associated with treatment conditions.

### Phenotypic and Survival Analysis

Whole-fly morphology was imaged using a stereomicroscope. Tumor-associated bloating was quantified by calculating the abdomen-to-head (A/H) area ratio, as previously described(Liu et al., 2025).

For lifespan analysis, adult flies were maintained on food containing either DMSO control or 30 µM Mitocurcumin, with survival scored every alternate day. Survival curves were plotted and compared using the log-rank (Mantel–Cox) test.

## Notes

### Competing Interest Statement

The authors have declared no competing interest.

