## Supplementary Information for "Mitocurcumin mediated redox disruption and metabolic rewiring induces tumor regression in *Drosophila* intestinal stem cell tumors"

**Supplementary figure legends:**

**Supplementary Figure S1. Conservation and stability of Mitocurcumin–TrxR interaction**

Chemical structures of curcumin and Mitocurcumin (A–B). Multiple sequence alignment of Drosophila TrxR1, mammalian TrxR2, and human TrxR1 (C). Ligand RMSD plots during molecular dynamics simulations (D, E). Protein backbone RMSD plots indicating structural stability. (F, G)

**Supplementary Figure S2. Expression profiling of anti-oxidant genes following Mitocurcumin treatment**

Schematic illustration of drug treatment regimen and tumor induction (A). Relative mRNA expression of antioxidant genes - *catalase* (B), *gtpx* (C) and *dSod2* (D) in adult midguts of MitoC treated tumor bearing flies compared to DMSO treated tumor bearing flies (B-D). RNA was isolated from 10 midguts per replicate (n = 3 biological replicates). Expression was normalized to *rp49*. Data represent mean ± SEM. Statistical significance was assessed using Student's t-test. p-values indicating statistical significance have been mentioned in the respective graphs.

**Supplementary Figure S3. Data normalization and pathway enrichment analysis for metabolomics**

Distribution of metabolite intensities before and after normalization, demonstrating improved comparability following $log_2$ transformation and Pareto scaling (A). Variable importance in projection (VIP) scores identifying metabolites contributing to group separation (B). Pathway enrichment analysis of significantly altered metabolites, showing enrichment in fatty acid metabolism, mitochondrial β-oxidation, TCA cycle, purine metabolism, and phospholipid biosynthesis (C–D). Metabolomic data were processed using Compound Discoverer and analyzed using MetaboAnalyst

Table 1. Molecular Docking and Binding Free Energy Parameters for Mitocurcumin-TrxR Complexes

| **Interaction Type** | **Drosophila TrxR1 (3DH9)** | **Mammalian TrxR1 (3QFA)** |
|---|---|---|
| **Glide Score (kcal/mol)** | **−9.02** | **−8.37** |
| **MM/GBSA ΔG(kcal/mol)** | **−28.92** | **−6.51** |
| **Hydrogen Bonds** | **Lys65**, **Glu333** | **Lys68**, **Lys69**, **Ser199**, **Asp334** |
| **Aromatic (π–π)** | **Tyr197** | **Tyr200** |
| **Hydrophobic** | Val58, Val60, Ile63, Ile196, Ala284, Ile285, Pro332, Leu334 | Val59, Val60, Val62, Ile65, Ile72, Leu112, Val201, Ile347 |
| **Polar** | Thr56, Ser177, Thr335 | Thr58, Thr343, Thr373 |
| **Charged (Basic)** | Lys26, Lys66, Arg223, Arg287, Lys307 | Arg166, Arg226, Arg293, Lys315, Arg351 |
| **Charged (Acidic)** | Asp117, Asp326 | Glu161, Glu163, Asp334 |

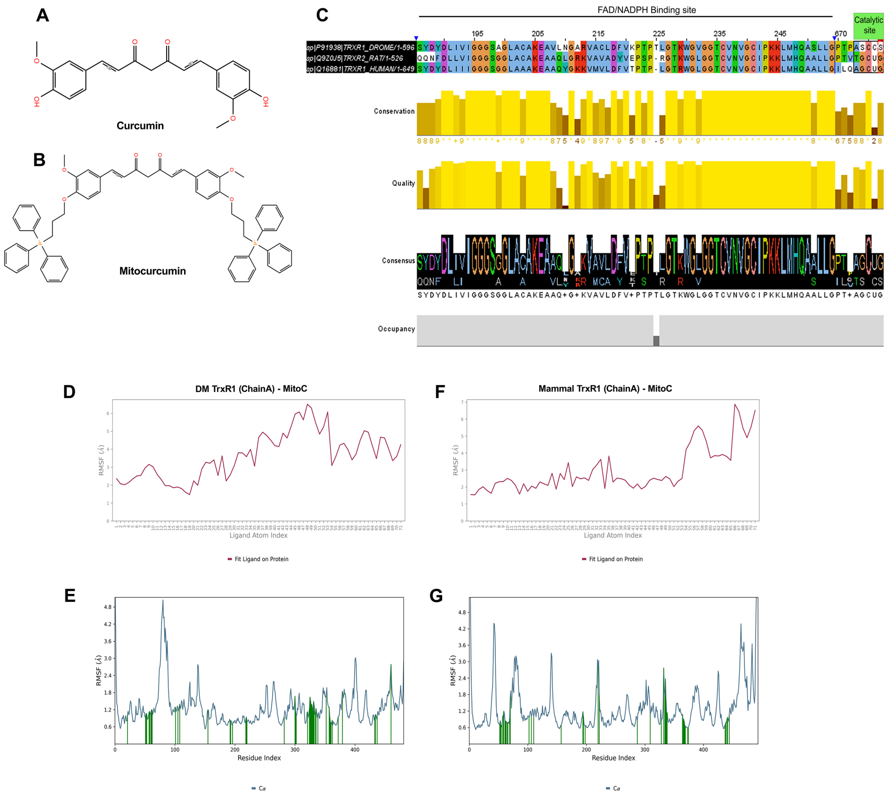

A
Curcumin
B
Mitocurcumin
C
FAD/NADPH Binding site
Catalytic site
sp|P91938|TRXR1_DROME/1-596
sp|Q9Z0J5|TRXR2_RAT/1-526
sp|Q16881|TRXR1_HUMAN/1-649
Conservation
Quality
Consensus
Occupancy
D
DM TrxR1 (ChainA) - MitoC
RMSF (Å)
Ligand Atom Index
Fit Ligand on Protein
F
Mammal TrxR1 (ChainA) - MitoC
RMSF (Å)
Ligand Atom Index
Fit Ligand on Protein
E
RMSF (Å)
Residue Index
Cα
G
RMSF (Å)
Residue Index
Cα

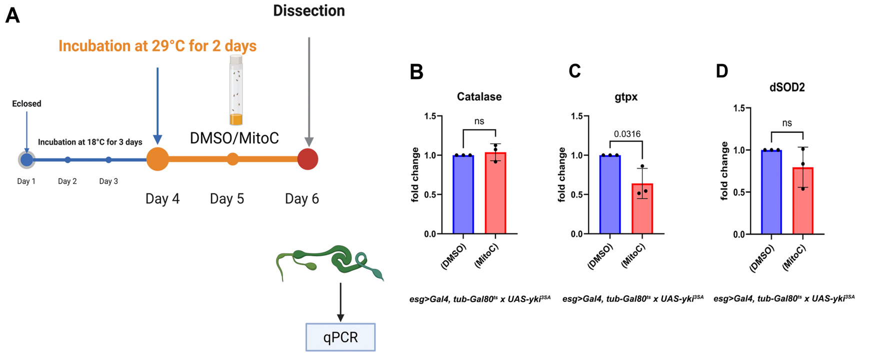
A
Dissection
Incubation at 29°C for 2 days
Eclosed
Incubation at 18°C for 3 days
DMSO/MitoC
Day 1
Day 2
Day 3
Day 4
Day 5
Day 6
qPCR
B
Catalase
ns
fold change
1.5
1.0
0.5
0.0
(DMSO)
(MitoC)
esg>Gal4, tub-Gal80ts x UAS-yki3SA
C
gtpx
0.0316
fold change
1.5
1.0
0.5
0.0
(DMSO)
(MitoC)
esg>Gal4, tub-Gal80ts x UAS-yki3SA
D
dSOD2
ns
fold change
1.5
1.0
0.5
0.0
(DMSO)
(MitoC)
esg>Gal4, tub-Gal80ts x UAS-yki3SA

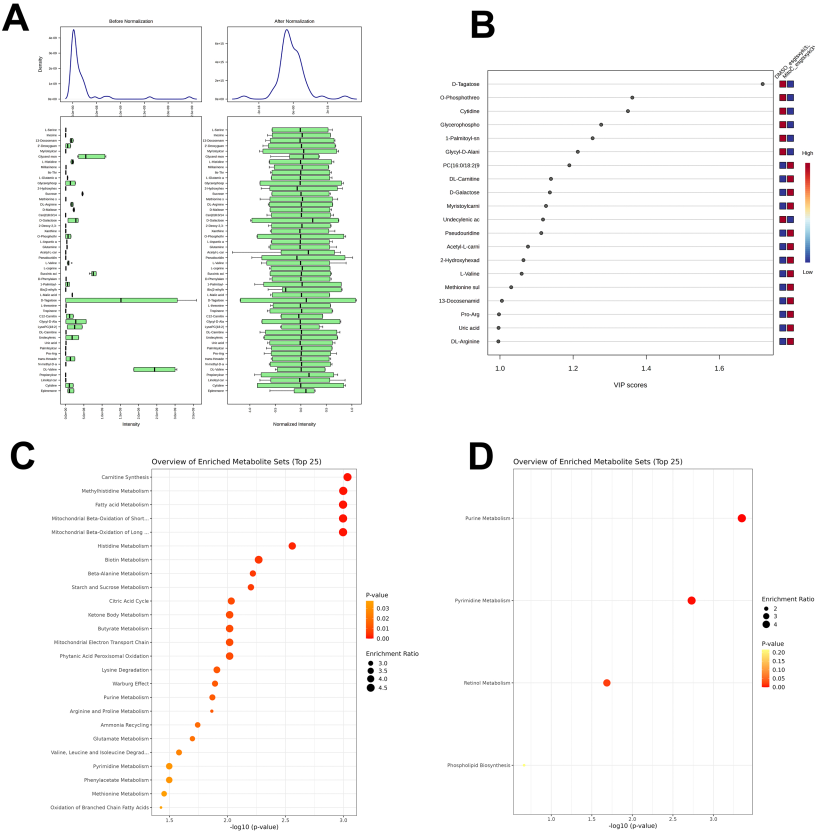
A
Before Normalization
After Normalization
Density
Intensity
Normalized Intensity
B
DMSO_esgtsxyki3
MitoC_esgtsxyki3
D-Tagatose
O-Phosphothreo
Cytidine
Glycerophospho
1-Palmitoyl-sn
Glycyl-D-Alani
PC(16:0/18:2(9
DL-Carnitine
D-Galactose
Myristoylcarni
Undecylenic ac
Pseudouridine
Acetyl-L-carni
2-Hydroxyhexad
L-Valine
Methionine sul
13-Docosenamid
Pro-Arg
Uric acid
DL-Arginine
High
Low
1.0
1.2
1.4
1.6
VIP scores
C
Overview of Enriched Metabolite Sets (Top 25)
Carnitine Synthesis
Methylhistidine Metabolism
Fatty acid Metabolism
Mitochondrial Beta-Oxidation of Short...
Mitochondrial Beta-Oxidation of Long ...
Histidine Metabolism
Biotin Metabolism
Beta-Alanine Metabolism
Starch and Sucrose Metabolism
Citric Acid Cycle
Ketone Body Metabolism
Butyrate Metabolism
Mitochondrial Electron Transport Chain
Phytanic Acid Peroxisomal Oxidation
Lysine Degradation
Warburg Effect
Purine Metabolism
Arginine and Proline Metabolism
Ammonia Recycling
Glutamate Metabolism
Valine, Leucine and Isoleucine Degrad...
Pyrimidine Metabolism
Phenylacetate Metabolism
Methionine Metabolism
Oxidation of Branched Chain Fatty Acids
1.5
2.0
2.5
3.0
-log10 (p-value)
P-value
0.03
0.02
0.01
0.00
Enrichment Ratio
3.0
3.5
4.0
4.5
D
Overview of Enriched Metabolite Sets (Top 25)
Purine Metabolism
Pyrimidine Metabolism
Retinol Metabolism
Phospholipid Biosynthesis
1.0
1.5
2.0
2.5
3.0
-log10 (p-value)
Enrichment Ratio
2
3
4
P-value
0.20
0.15
0.10
0.05
0.00

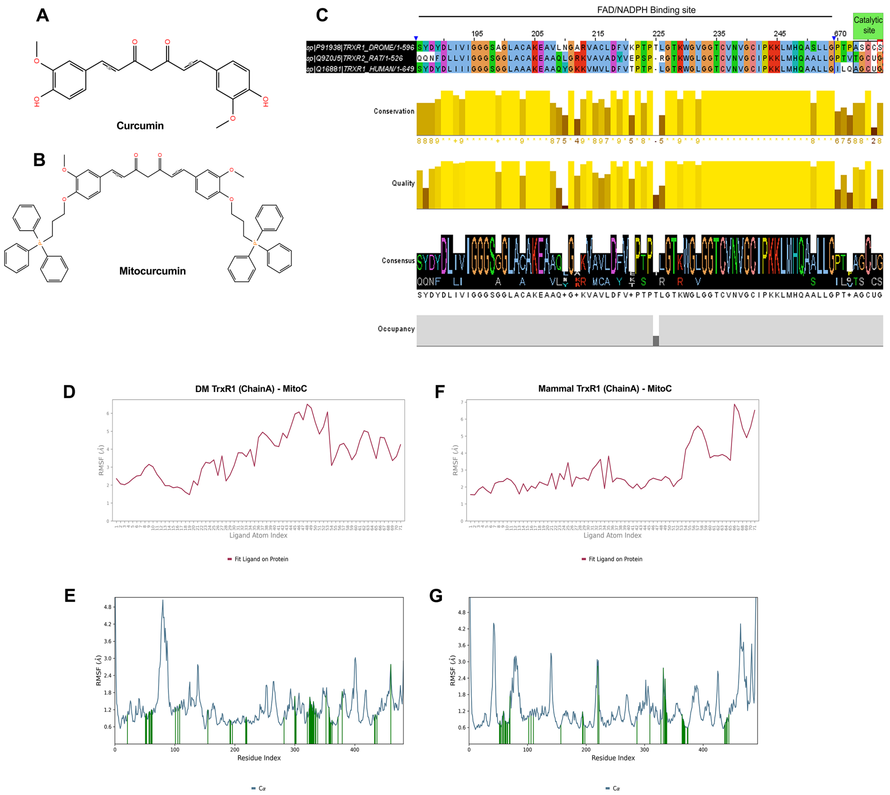
A
Curcumin
B
Mitocurcumin
C
FAD/NADPH Binding site
Catalytic site
195
205
215
225
235
245
670
sp|P91938|TRXR1_DROME/1-596
sp|Q9Z0J5|TRXR2_RAT/1-526
sp|Q16881|TRXR1_HUMAN/1-649
Conservation
Quality
Consensus
SYDYDLIVIGGGSGGLACAKEAAQ+G+KVAVLDFV+PTPTLGTKWGLGGTCVNVGCIPKKLMHQAALLGPT+AGCUG
Occupancy
D
DM TrxR1 (ChainA) - MitoC
RMSF (Å)
Ligand Atom Index
Fit Ligand on Protein
F
Mammal TrxR1 (ChainA) - MitoC
RMSF (Å)
Ligand Atom Index
Fit Ligand on Protein
E
RMSF (Å)
Residue Index
Cα
G
RMSF (Å)
Residue Index
Cα

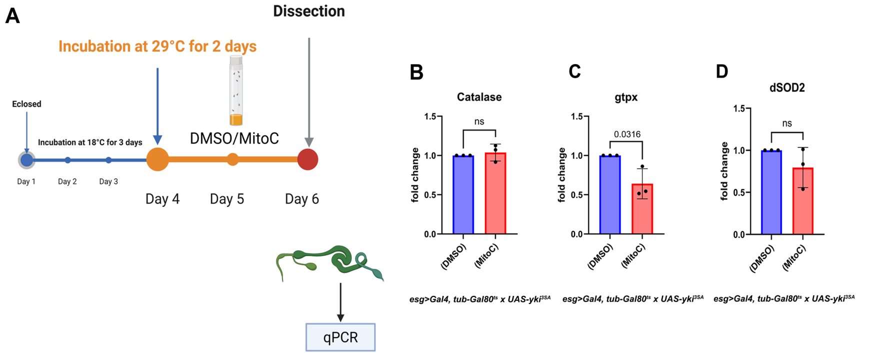
A
Dissection
Incubation at 29°C for 2 days
Eclosed
Incubation at 18°C for 3 days
DMSO/MitoC
Day 1
Day 2
Day 3
Day 4
Day 5
Day 6
qPCR
B
Catalase
ns
fold change
1.5
1.0
0.5
0.0
(DMSO)
(MitoC)
C
gtpx
0.0316
fold change
1.5
1.0
0.5
0.0
(DMSO)
(MitoC)
D
dSOD2
ns
fold change
1.5
1.0
0.5
0.0
(DMSO)
(MitoC)
esg>Gal4, tub-Gal80ts x UAS-yki3SA
esg>Gal4, tub-Gal80ts x UAS-yki3SA
esg>Gal4, tub-Gal80ts x UAS-yki3SA

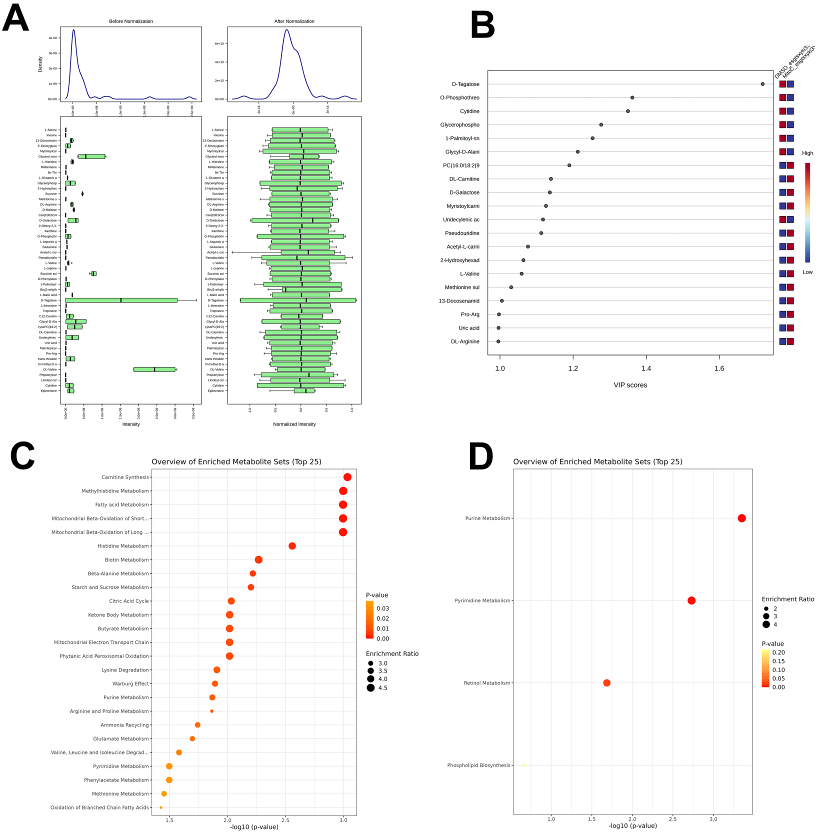
A
Before Normalization
After Normalization
Density
Intensity
Normalized Intensity
B
DMSO_esgtsxyki3
MitoC_esgtsxyki3
D-Tagatose
O-Phosphothreo
Cytidine
Glycerophospho
1-Palmitoyl-sn
Glycyl-D-Alani
PC(16:0/18:2(9
DL-Carnitine
D-Galactose
Myristoylcarni
Undecylenic ac
Pseudouridine
Acetyl-L-carni
2-Hydroxyhexad
L-Valine
Methionine sul
13-Docosenamid
Pro-Arg
Uric acid
DL-Arginine
High
Low
1.0
1.2
1.4
1.6
VIP scores
C
Overview of Enriched Metabolite Sets (Top 25)
Carnitine Synthesis
Methylhistidine Metabolism
Fatty acid Metabolism
Mitochondrial Beta-Oxidation of Short...
Mitochondrial Beta-Oxidation of Long ...
Histidine Metabolism
Biotin Metabolism
Beta-Alanine Metabolism
Starch and Sucrose Metabolism
Citric Acid Cycle
Ketone Body Metabolism
Butyrate Metabolism
Mitochondrial Electron Transport Chain
Phytanic Acid Peroxisomal Oxidation
Lysine Degradation
Warburg Effect
Purine Metabolism
Arginine and Proline Metabolism
Ammonia Recycling
Glutamate Metabolism
Valine, Leucine and Isoleucine Degrad...
Pyrimidine Metabolism
Phenylacetate Metabolism
Methionine Metabolism
Oxidation of Branched Chain Fatty Acids
1.5
2.0
2.5
3.0
-log10 (p-value)
P-value
0.03
0.02
0.01
0.00
Enrichment Ratio
3.0
3.5
4.0
4.5
D
Overview of Enriched Metabolite Sets (Top 25)
Purine Metabolism
Pyrimidine Metabolism
Retinol Metabolism
Phospholipid Biosynthesis
1.0
1.5
2.0
2.5
3.0
-log10 (p-value)
Enrichment Ratio
2
3
4
P-value
0.20
0.15
0.10
0.05
0.00
